# Beyond homogeneity: Charting the landscape of heterogeneity in psychiatric electroencephalography

**DOI:** 10.1101/2024.03.04.583393

**Authors:** Aida Ebadi, Sahar Allouch, Ahmad Mheich, Judie Tabbal, Aya Kabbara, Gabriel Robert, Aline Lefebvre, Anton Iftimovici, Borja Rodríguez-Herreros, Nadia Chabane, Mahmoud Hassan

**Affiliations:** MINDIG, F-35000, Rennes, France; Service des Troubles du Spectre de l’Autisme et apparentés, Département de psychiatrie, Lausanne University Hospital (CHUV), Lausanne, Switzerland; Faculty of science, Lebanese International University, Tripoli, Lebanon; Pôle de Psychiatrie Hospitalo-Universitaire de Psychiatrie Adulte, Centre Hospitalier Guillaume Régnier, Rennes, France; U1228 Empenn UMR 6074 IRISA, Rennes, France; Paris Saclay University, Neurospin, CEA Saclay, Service Avis et Expertise TND, Fondation Vallée; Université Paris Cité, Institute of Psychiatry and Neuroscience of Paris (IPNP), INSERM U1266, Team “Pathophysiology of psychiatric disorders”, GDR 3557-Institut de Psychiatrie, Paris, France; GHU Paris Psychiatrie et Neurosciences, Pôle hospitalo-universitaire d’évaluation, prévention, et innovation thérapeutique (PEPIT), Paris, France; School of Science and Engineering, Reykjavik University, Reykjavik, Iceland

**Author notes:** Corresponding authors: Sahar Allouch and Mahmoud Hassan.

## Abstract

Electroencephalography (EEG) has been thoroughly studied for decades in psychiatry research. Yet its integration into clinical practice as a diagnostic/prognostic tool remains unachieved. We hypothesize that a key reason is the underlying patient’s heterogeneity, overlooked in psychiatric EEG research relying on a case-control approach. We combine HD-EEG with normative modeling to quantify this heterogeneity using two well-established and extensively investigated EEG characteristics -spectral power and functional connectivity-across a cohort of 1674 patients with attention-deficit/hyperactivity disorder, autism spectrum disorder, learning disorder, or anxiety, and 560 matched controls. Normative models showed that deviations from population norms among patients were highly heterogeneous and frequency-dependent. Deviation spatial overlap across patients did not exceed 40% and 24% for spectral and connectivity, respectively. Considering individual deviations in patients has significantly enhanced comparative analysis, and the identification of patient-specific markers has demonstrated a correlation with clinical assessments, representing a crucial step towards attaining precision psychiatry through EEG.

## Introduction

Electroencephalography (EEG) has been extensively studied in psychiatry research to identify electrophysiological correlates of various disorders (Loo and Makeig 2012; Olbrich and Arns 2013; de Aguiar Neto and Rosa 2019; Wang et al. 2013), disease severity (Livint Popa et al. 2020), subtypes (Zhang et al. 2021; Slater et al. 2022; Clarke et al. 2001), and treatment response (Widge et al. 2019; Watts et al. 2022; Wu et al. 2020; Rolle et al. 2020). The non-invasive nature of EEG, its cost-effectiveness, and its capacity to capture rapid spontaneous brain activity make it a highly appealing tool for clinical psychiatry. Nonetheless, despite considerable research efforts, the pathophysiological mechanisms of psychiatric disorders are still poorly understood and the progress in establishing clinically applicable EEG-based biomarkers has fallen short of expectations. This is evidenced by the pronounced inconsistencies observed across studies (Newson and Thiagarajan 2019; Neo et al. 2023; González-Madruga, Staginnus, and Fairchild 2022; Cortese et al. 2021; Miljevic et al. 2023).

A key factor contributing to this issue is the reliance, in EEG research, on statistical designs that fail to account for the inherent heterogeneous nature of psychiatric disorders (Feczko et al. 2019; Marquand et al. 2019). Typically, these designs rely on comparisons of average differences between groups (e.g. patient versus control or treatment versus placebo), presupposing uniformity within groups, as well as distinct separations between cases and healthy controls (Verdi et al. 2021; Marquand et al. 2016, 2019; Feczko et al. 2019). However, such assumptions significantly misrepresent the reality of psychiatric disorders, which exhibit profound heterogeneity and overlap in symptoms, severity levels, developmental course, biological underpinnings, and response to treatment (Segal et al. 2023).

To more effectively capture the nuanced heterogeneity of psychiatric disorders, research methodologies should go beyond conventional case-control paradigms to more sophisticated statistical techniques that can accommodate the variability within and across patient and healthy populations (Marquand et al. 2019; Verdi et al. 2021). An emerging powerful approach in neuroimaging is normative modeling (NM), which involves estimating normative trajectories of a reference population and assessing the degree to which individuals deviate from these norms (Verdi et al. 2021; Marquand et al. 2016, 2019). A well-known example of NM is pediatric growth charts, by which a child’s height and weight are compared to those of children of the same age and sex (Cole 2012). Recent MRI studies combined with NM have endeavored to chart analogous trajectories for brain phenotypes (e.g., grey and white matter volumes, mean cortical thickness, total surface area, etc.) to map lifespan age-related changes in brain structure (Bethlehem et al. 2022; Rutherford, Fraza, et al. 2022; Rutherford et al. 2023) and characterize structural heterogeneity in psychiatric disorders (Segal et al. 2023) such as ADHD (Wolfers et al. 2020), Autism (Zabihi et al. 2020), Schizophrenia and Bipolar disorder (Wolfers et al. 2018), as well as in neurodegenerative disease as Alzheihmer’s disease (Verdi et al. 2023). Additionally, there has been a development of normative models for functional MRI, although to a lesser extent (Rutherford, Kia, et al. 2022; Sun et al. 2023). However, to our knowledge, similar research focusing on the electrophysiological aspects of psychiatric disorders and the underlying substrates of heterogeneity remains unexplored.

Here, we address this gap by combining HD-EEG with NM to elucidate the electrophysiological heterogeneity of psychiatric disorders. We perform our investigation at two levels, the scalp (the level of electrode) and the sources (the level of cortical brain sources), thanks to the availability of high channel density. We develop an end-to-end NM framework of a normative HD-EEG cohort comprising 560 healthy subjects (age 5-18 yo), see Fig. 1a for illustrative explanation. We then systematically quantify the heterogeneity observed in the HD-EEG power spectrum and functional connectivity among a clinical group of 1674 age-matched patients diagnosed with autism ASD, Attention deficit hyperactivity disorder (ADHD), anxiety (ANX), and learning disorders (LD), Fig. 1b, 1c. Inspired by MRI-based studies, we hypothesize that psychiatric disorders exhibit substantial electrophysiological heterogeneity and that individual patient deviations are likely to demonstrate low spatial homogeneity across both channels and cortical regions.

**Fig. 1.**
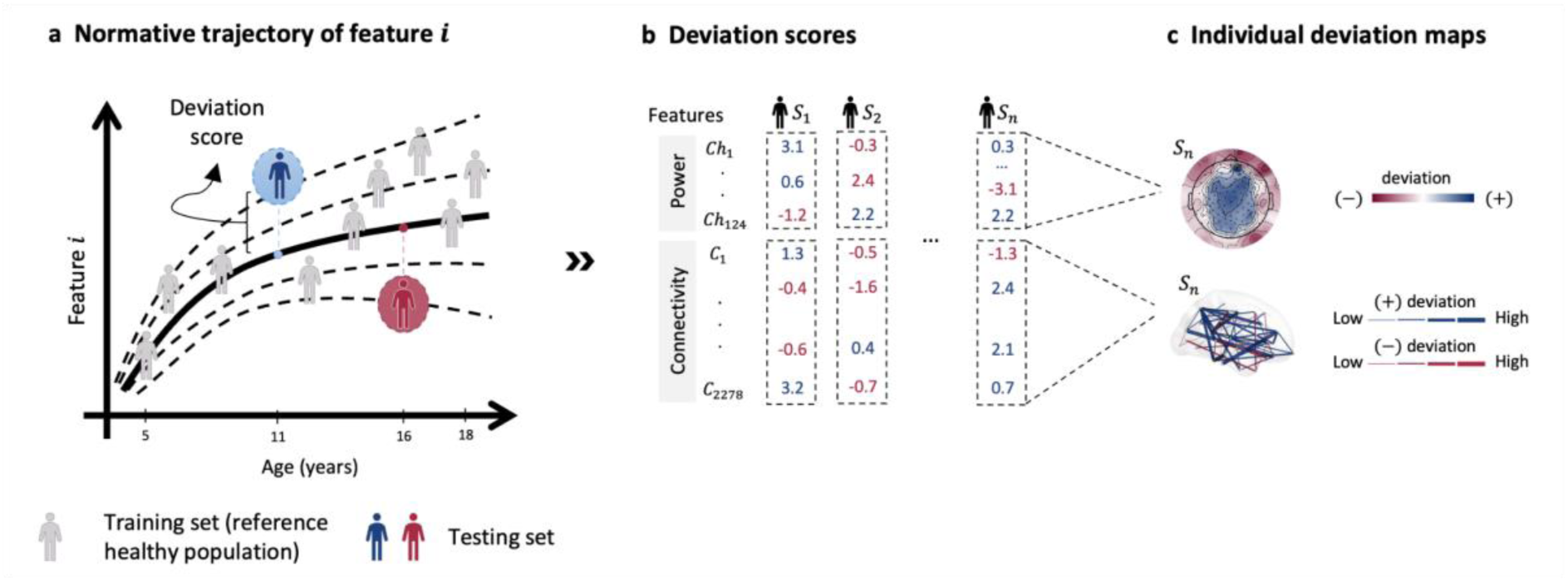
Normative modeling of electrophysiological features across age and individual deviation assessment. (a) Illustration of the normative trajectory of a given feature across different ages, with the solid line representing the median trajectory derived from a healthy reference population (training set). (b) Tabulation of deviation scores for individual subjects across multiple features including power spectral densities and functional connectivity. Red and blue values denote negative and positive deviations (i.e., below/above the median), respectively. (c) Individual deviation maps showing spectral (upper panel) and network-based (lower panel) deviations.

## Results

### Data description

High-density (128 channels) resting-state EEG data was collected from subjects aged between 5 and 18 years old across multiple datasets (<u>Methods</u>). Models were trained on 448 (52% M) healthy controls (HC) and 112 (55% M) healthy subjects were held out as a comparison group against the clinical cohort. The clinical groups comprised 576 subjects diagnosed with ASD (52% M), 650 with ADHD (27% M), 216 with Anxiety disorders (52% M), and 232 with Learning disorders (46% M).

### Normative modeling

EEG power spectra (i.e., EEG power in the predefined frequency band: delta, theta, alpha, beta, and gamma) have been the most used EEG features in psychiatry (Neo et al. 2023). More recently and with the availability of high-density EEG, source-space functional connectivity analysis has emerged as a highly promising approach, establishing a framework for the electrophysiological circuit-level differentiation of healthy and diseased brains (Hassan and Wendling 2018). Here, we have developed models for the normative trajectories of these two well-established sets of features and assessed the heterogeneity in psychiatric disorders based on subject deviations from the normative trajectories. For each feature, we trained a Generalized Additive Model for Location, Scale, and Shape (GAMLSS) on a healthy reference population. For the spectral features, a GAMLSS was fitted to the relative power (Welch’s method) of each channel and each frequency band. Functional connectivity was estimated between 68 predefined brain regions from the Desikan-Killiany atlas using Amplitude Envelope Correlation -AEC-(Brookes et al. 2011; Hipp et al. 2012), corrected for source leakage using an orthogonalization approach (Brookes, Woolrich, and Barnes 2012). A GAMLSS was fitted to the connectivity values at each connection and each frequency band.

Subsequently, subjects with psychiatric disorders (ASD, ADHD, ANX, and LD) along with an HC(test) group were projected on these models to calculate their deviation scores (*z*-scores, *z*). These deviations can be negative (lower than normative values) or positive (higher than normative values), Fig. 2a1, 2b1. Extreme deviations were defined as |*z|*> 2 (Bethlehem et al. 2022; Rutherford, Kia, et al. 2022; Rutherford et al. 2023; Segal et al. 2023; Wolfers et al. 2020, 2018; Zabihi et al. 2020; Verdi et al. 2023), Fig. 2a2, 2b2.

**Fig. 2.**
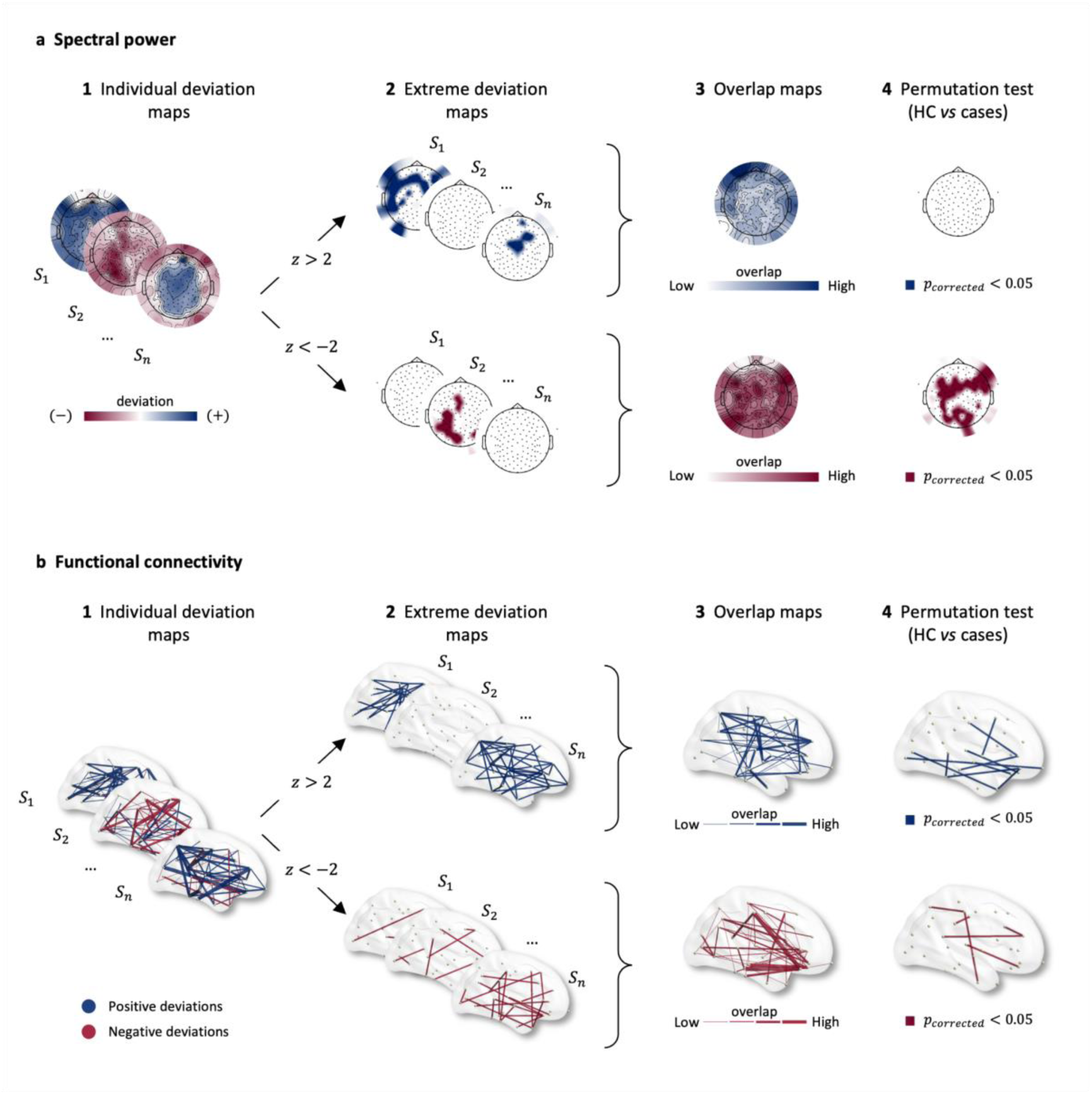
Overview of the quantification of heterogeneity in spectral and connectivity features based on normative model-inferred deviation scores. **(a1)** individualized deviation maps scores for spectral power, (**a2)** extreme deviation maps displaying both positive (top) and negative (bottom) deviations, identified for deviation scores >2 and <-2, respectively, **(a3)** spatial overlap maps, calculated as the percentage of subjects showing extreme deviation at each channel, among those with at least one extreme deviation, to identify common areas of deviation. **(a4)** group-based permutation tests to evaluate group differences (HC(test) *vs* clinical group) in channel-level overlap (*p<0.05*, FDR corrected) **(b1-b4)** same methodological approach applied for functional connectivity.

First, we computed the percentage of subjects with at least one extremely deviated channel/connection, as well as the number of extremely deviated channels/connections per subject in each group. Next, we evaluated the spatial overlap of extreme deviations by computing the percentage of extremely deviated subjects per channel/connection within each group. (Fig. 2a3, 2b3). A group-based permutation test (Segal et al. 2023) was used to compare clinical groups and HCs overlap maps (all results were corrected for multiple comparisons using FDR), Fig. 2a4, 2b4.

The distribution of the number of extremely deviated channels/connections across groups is depicted in Fig. S19-S28 in Supplementary Materials. Range and median values are reported in Table S6-S15. The number of negatively deviated channels per subject in all clinical groups were significantly different as compared to HC(test) in delta band (*p*<0.03, Mann-Whitney test). For the positive deviations, all groups showed significant differences compared to HC(test) at alpha and beta bands (*p<0.01*). Regarding functional connectivity features, ADHD and ASD showed significantly fewer numbers of negative extreme deviations compared to HCs, particularly in the beta and gamma bands (*p<0.01*). Conversely, ANX and LD demonstrate significant differences only in the alpha band (*p<0.05*). However, connections displaying positive deviations are significant across all clinical groups and frequency bands (*p<0.01*). Interestingly, while we observed significant differences in most cases, our results did not reveal a uniform trend. In some instances, clinical cohorts exhibited higher numbers of extreme deviations compared to the HC group; conversely, in other instances, the HC group showed more extreme deviations than clinical groups.

### Spectral heterogeneity

The percentages of subjects having at least one negative deviation were relatively low with the highest values occurring in the theta (ASD: 44%), alpha (ADHD: 29%, LD: 25%, ANX: %23), and beta (HC(test): %35) bands (These percentages are detailed across all frequency bands and disorders in Table S6-S10 in the Supplementary Materials). Thus, a substantial proportion of participants exhibit significant similarities with the healthy control (HC) group. Then, we quantified the spatial overlap of these extreme deviations across subjects. Fig. 3a illustrates examples of the overlap maps across groups (detailed results figures can be found in Supplementary Materials, Fig S29-S33). Interestingly, the overlap values were low and did not exceed 40% (delta: ADHD: 31%, ANX: 32%; beta: ASD: 28%, HC(test): 31%; gamma: LD: 40%) reflecting an inconsistency in the spatial location of extreme deviations. Notably, LD and ADHD exhibited the highest consistency (i.e., overlap) across subjects, mainly in the delta band. Overall, similar results were obtained for positive deviations. The ASD group exhibited the highest percentage of subjects having at least one extreme deviation (41% in theta band) and the spatial overlap that did not exceed 40% (HC(test), alpha).

**Fig. 3.**
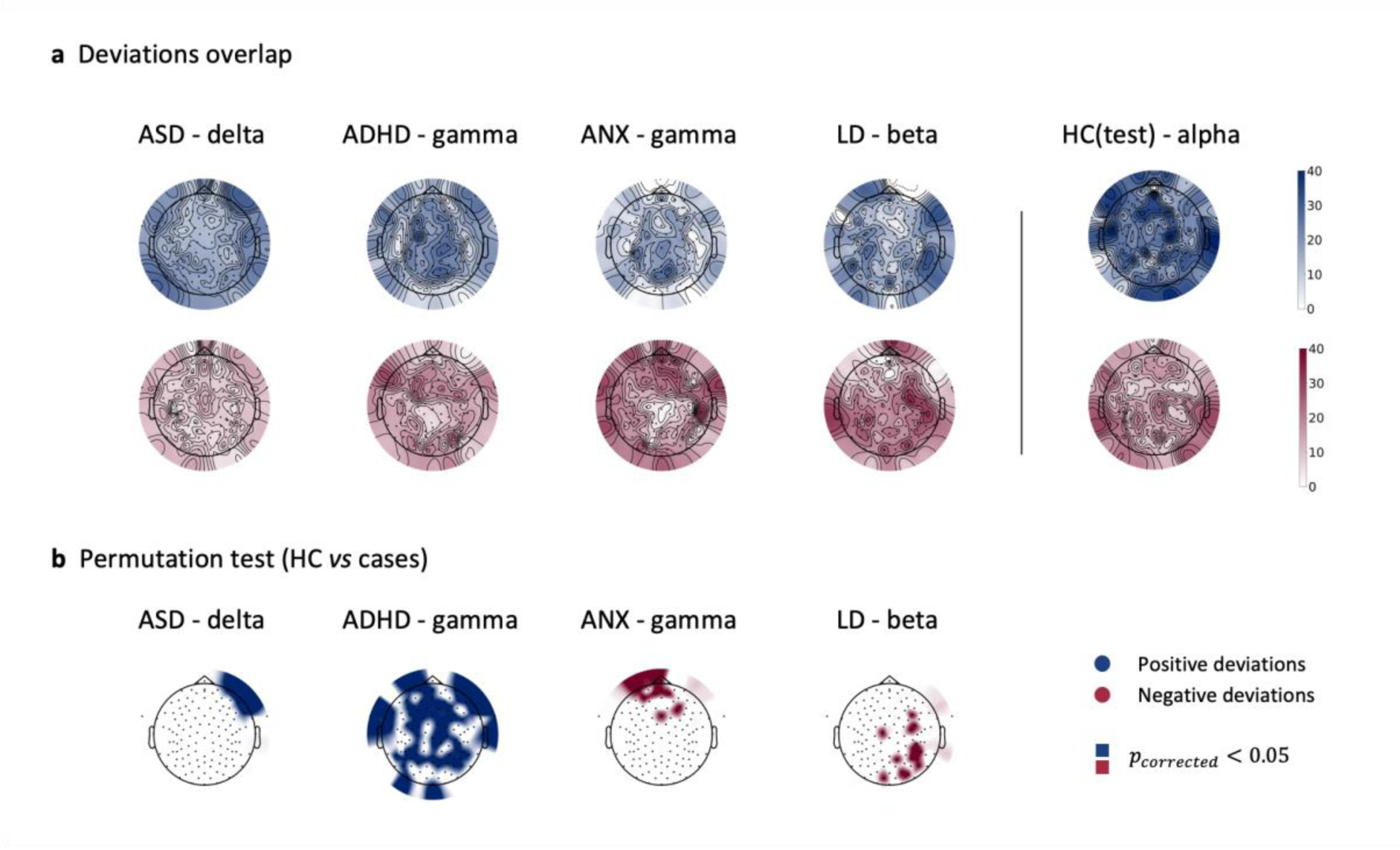
Spectral features heterogeneity. **(a)** Overlap maps of deviation scores for clinical groups and the held-out healthy control group (HC(test)), illustrating areas of common deviation. (**b)** channels showing significant differences between HC(test) and clinical groups, determined through group-based permutation tests (*p<0.05*, *FDR corrected*).

To investigate if the obtained overlap may differentiate patients from HC, we compared the channel-wise overlap maps between clinical groups and healthy controls. The result of group-based permutations (see <u>Methods</u>) varies greatly between frequency bands (Fig. 3b). Out of 124 channels (remained after data preprocessing), the highest number of channels showing significant difference with HC(test) for negative extreme deviation overlap was 41 (delta: ADHD: 26; beta: ASD: 19, LD: 41; ANX: 22; *uncorrected*). Only 15 channels survive FDR correction for LD in beta band mainly in the occipital-parietal region. For anxiety in the beta band, none survive the FDR correction, instead in the gamma band 11 channels remain mostly in the frontal area (Fig. 3b). For positive extreme deviations, ADHD has the highest number of significant channels in gamma (delta: ANX: 50; alpha: ASD: 57; gamma: ADHD: 86, LD: 63, *uncorrected*), of which 74 channels remain after FDR correction. For the ASD group after FDR correction there are no significant channels in the alpha band, only 3 channels in the frontal region survive in the delta band (Fig. 3b). This low overlap and weak differentiation between cases and HC confirm the absence of consistent characteristic patterns among a whole group. Instead, the patterns seem to be subject-specific. All the results for cases and frequency bands are presented in the supplementary materials (Figure S29 to S33).

### Heterogeneity at the level of functional connectivity

Next, we quantified the heterogeneity at the level of functional brain connections. Unlike the findings from spectral analysis, we observed an increase in the number of subjects exhibiting at least one negative/positive extreme deviation. Detailed percentages across all frequency bands and disorders are provided in Table S11-S15 in the Supplementary Materials. For negative extreme deviations, the highest observed percentage was 87% (delta: HC(test): 87%, ASD: 71%; theta: ADHD: 82%, ANX: 83%, LD: 82%). Among subjects with at least one extreme deviation, the spatial overlap of these deviations was notably low, not exceeding 14% (alpha: HC(test): 14%, ASD: 9%, ADHD: 10%, LD: 10%, gamma: ANX: 12%), Fig. 4b. For positive deviations, the percentage of subjects with at least one extremely deviant connection is modestly lower (the highest value (73%) obtained for ASD in beta band). Conversely, the within-groups overlap percentages slightly increased (delta: ADHD: 15%; theta: ANX: 19%; alpha: HC(test): 23%, beta: LD: 24%, gamma: ASD: 21%) with LD exhibiting the highest overlap values, Fig. 4a.

**Fig. 4.**
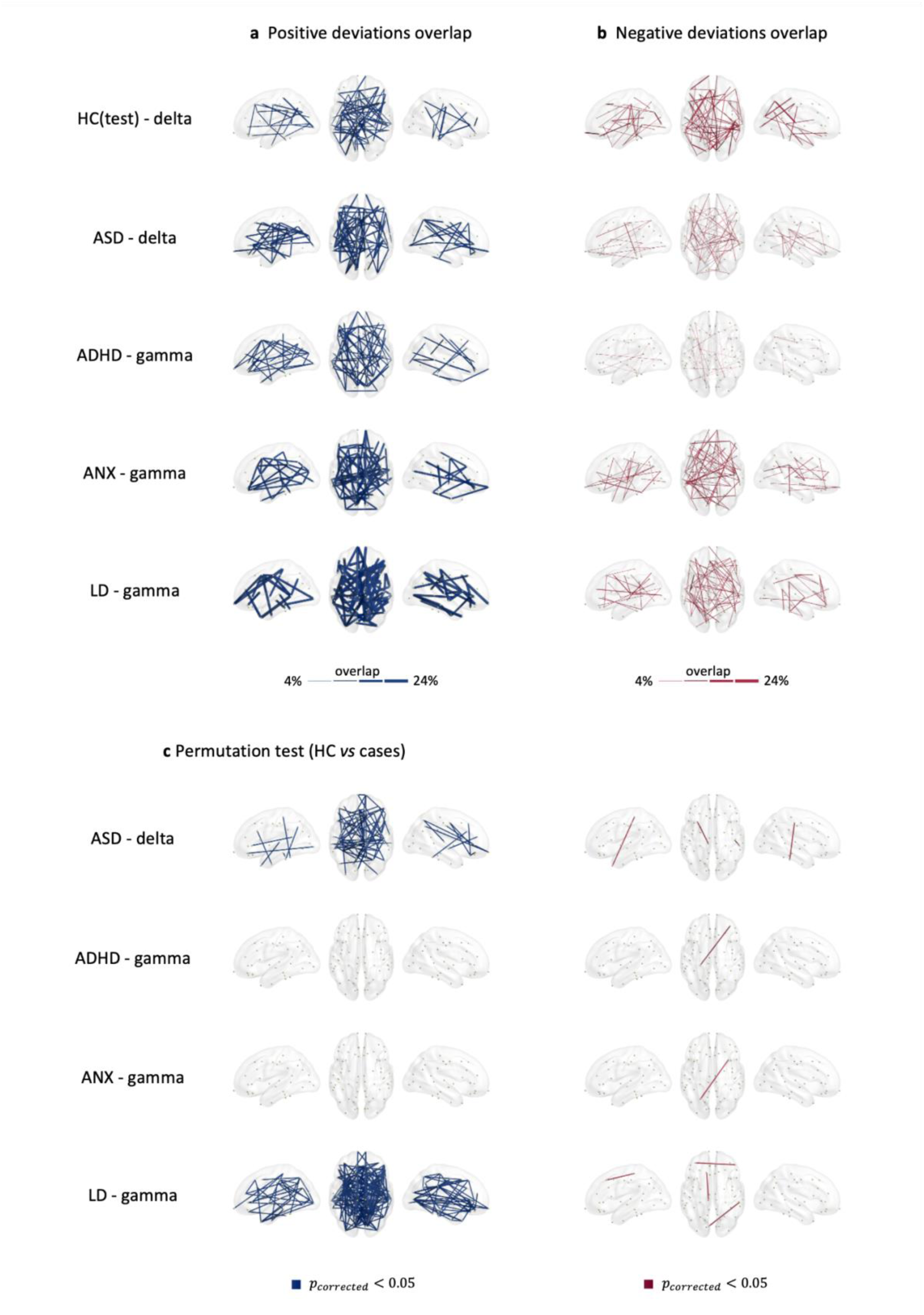
Functional connectivity heterogeneity. Overlap maps of **(a)** positive and **(b)** negative deviation scores for clinical groups and the held-out healthy control group (HC(test)) within the different frequency bands, illustrating areas of common deviation among patients (with only the highest 3% overlap values being plotted for visualization purposes). **(c-d)** functional connections showing significant differences between HC(test) and clinical groups at the different frequency bands, determined through group-based permutation tests (*p<0.05*, *FDR corrected*).

Significant differences, using group-based permutation tests corrected using FDR, in overlap maps between clinical groups and healthy controls are shown in Fig. 4c and Fig S34-S38 in Supplementary Materials. The variation is pronounced across frequency bands. Compared to connections with negative deviations, a higher number of connections with positive deviations were found to be significant.

### Deviation scores as patient-specific markers for comparative analysis and correlation with clinical assessment

Having assessed the heterogeneity among patients, we sought potential solutions by leveraging NMs to derive EEG-markers tailored to individual patients. To explore this, we performed two proof-of-concept analyses. First, we explored whether the metrics derived from NM could outperform classical features in differentiating between healthy controls and patient groups. Using a network-based statistics -NBS-approach (permutation test, n=5000) (Zalesky, Fornito, and Bullmore 2010), we compared the clinical groups to the HC using either the original functional connectivity matrices (the original features) or the deviation score (z-scores) matrices. Interestingly, FC features did not show any significant differences between HC *vs.* cases. However, using *z*-scores, significant differences were found between groups at different frequency bands. Examples of these differences are presented in Fig. 5a for ASD at alpha (*p*=0.0002, 91 edges) and beta (*p*=0.007, 12 edges) bands and ADHD at delta band (*p*<0.0001, 94 edges).

**Fig. 5.**
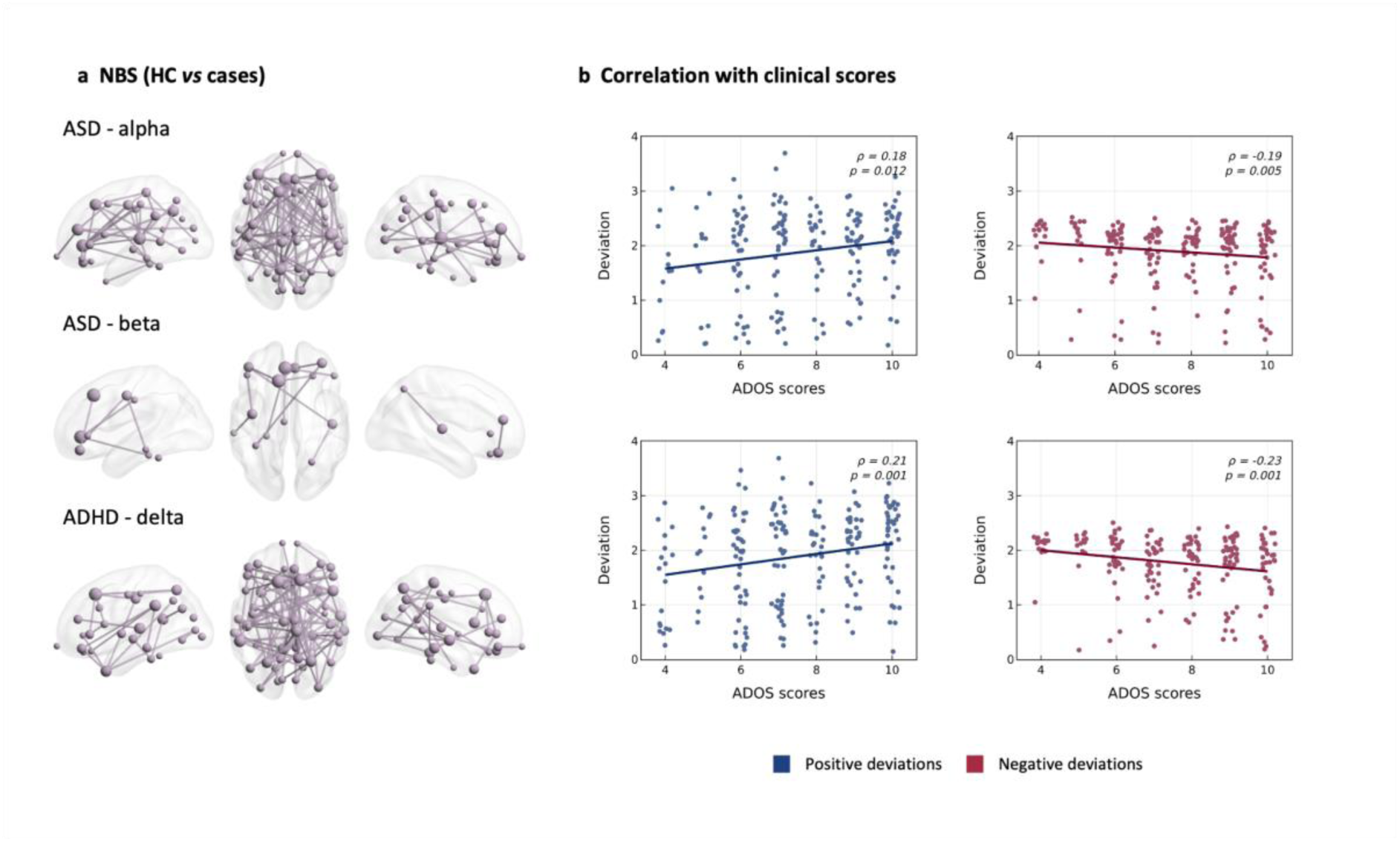
Leveraging deviation scores for group-based analysis and correlation with clinical assessments. (a) Identification of significant network patterns that differentiate case groups (ASD and ADHD) from the healthy control (HC) group, as revealed by network-based statistics (NBS). (b) Correlation between ASD subjects’ global deviation scores and clinical assessment scores (total ADOS) in alpha and gamma bands.

Second, we explored the potential of NM to provide a framework for generating subject-specific markers that may correlate with clinical assessments of patients. For this purpose, we calculated a global deviation score for each subject, defined as the average *z* values of the extremely deviated connections. This score was then correlated with the patients’ clinical assessments. Fig. 5b shows a significant, but relatively low, Spearman correlation (average *ρ*=0.2, *p*<0.05) between total ADOS scores (a standard clinical assessment in ASD) and both positive and negative patient-specific global deviation scores. As ADOS scores increased, positive global deviations also increased, while negative global deviations decreased. Global deviations derived from the spectral models showed no correlation with clinical scores. These findings are a first step into developing EEG-based patient-specific markers that can be used for objectively quantifying personalized treatment such as medication or neurostimulation.

## Discussion

We combined HD-EEG with normative modeling to characterize the heterogeneity in spectral power and functional connectivity among patients with psychiatric disorders. We showed highly heterogeneous alterations with deviation spatial overlap across patients that did not exceed 40% and 24% for spectral power and connectivity, respectively. Our results challenge the prevailing reliance on a case-control approach in psychiatric EEG studies, emphasizing the importance of recognizing individual variability. The pronounced heterogeneity we observed across the conditions studied suggests that assuming within-group homogeneity, as implicitly done in prior psychiatric EEG research, oversimplifies the intricate neurophysiological signatures associated with psychiatric disorders. We finally showed that through the consideration of the individual patient variability, the enhancement of comparative analysis has been substantial. The identification of patient-specific markers has demonstrated a correlation with clinical assessments, representing a crucial advancement in the pursuit of EEG precision psychiatry.

### Heterogeneity in EEG spectral and connectivity features

EEG spectral power has long dominated EEG clinical research in general and specifically in psychiatry (Neo et al. 2023). Investigating changes in the power of predefined frequency bands has been one of the standards in EEG research for determining changes between healthy controls and patient groups. However, research outcomes are usually inconsistent across studies (Neo et al. 2023; Newson and Thiagarajan 2019). Moreover, changes in EEG power do not characterize a single disorder but rather show significant overlap across various psychiatric conditions (Newson and Thiagarajan 2019). For instance, an increase in power in the lower frequency bands (delta and theta) and a decrease across higher frequencies (alpha, beta, and gamma) represent a dominant pattern of change across several disorders, including ADHD, schizophrenia, and OCD (Newson and Thiagarajan 2019). Furthermore, a significant number of conditions, including PTSD, addiction, and autism, do not exhibit a consistent pattern of spectral change in any specific direction (Newson and Thiagarajan 2019).

Functional connectivity research represents an emerging framework that is not as established as spectral analysis. Despite its novelty, there is a significant body of studies aimed at characterizing functional alterations associated with psychiatric disorders. Nevertheless, inconsistencies in the results have been observed (Miljevic et al. 2023). For instance, meta-analyses of resting-state functional connectivity in ADHD found no spatial convergence across studies (González-Madruga, Staginnus, and Fairchild 2022; Cortese et al. 2021). Similar findings were observed in (Samea et al. 2019) where no significant convergent functional alterations in children/adolescents with ADHD in their main meta-analysis comprising 1914 unique participants from 96 studies.

The observed inconsistency in results can be, in part, justified by the lack of a standardized methodology for EEG data acquisition and analysis (variations in electrode configurations, task paradigms, signal processing techniques, etc…) and small sample sizes compromising the generalizability of the findings. However, even with efforts to control for these methodological variables, achieving consistent results remains challenging. A principal source of this variability is the inherent heterogeneity among patient populations in psychiatric disorders (Feczko et al. 2019; Marquand et al. 2016), which may introduce confounding factors not accounted for in group-level analyses. The heterogeneity we observed here in psychiatric EEG is in line with recent studies using MRI (structural and functional) with NM in psychiatry. Indeed, it was shown that patient-specific deviations from population expectations for regional gray matter volume were highly heterogeneous, affecting the same area in <7% of people with the same diagnosis (Segal et al. 2023). Our results at the channel/connection level showed higher consistency than these structural MRI-based studies. Our results are however comparable with the results obtained by these studies when looking for overlaps at network/circuit level (∼40 to 50%).

We believe that the demonstrated heterogeneity is the primary factor hindering the development of EEG-based biomarkers. The challenges associated with group-level analysis manifest at different levels. At the diagnostic level, patients are often classified into distinct, clearly defined groups, presupposing homogeneity within each group. At the treatment level, a ’*one-size-fits-all*’ strategy is often adopted, applying the same treatment protocols to all patients without considering individual heterogeneity. To address this issue, it is crucial to develop patient-specific electrophysiological biomarkers that aim to 1) accurately diagnose disease conditions, 2) monitor and predict disease progression, and 3) guide patients in choosing therapeutic options tailored to their individual risk factors.

### Beyond heterogeneity mapping

Here, by leveraging subject-level inferences from normative models, we have mapped the heterogeneity within psychiatric conditions and among healthy populations. However, the utility of normative models extends beyond merely elucidating sample heterogeneity. Deviation scores obtained from these models also can be used as inputs for downstream analyses. Instead of using the raw features (i.e. relative power, functional connectivity, etc.), group average, classification, and prediction analyses can be run using the deviation scores inferred from the models that can serve as inputs. For instance, (Rutherford et al. 2023) found minor (regression) to strong (group difference testing) advantages of using deviation scores over raw features. We have indeed tested this in the current paper and showed that NM can indeed improve group-level comparison between HC and cases. As such, normative models can contribute to the development of more personalized electrophysiological approaches.

Moreover, one of the main aims of combining EEG and normative modeling, in addition to deciphering heterogeneity, is to develop a patient-specific marker that can be clinically useful. We analyzed the clinical correlates of extreme deviations and showed associations between individual-specific deviations and their clinical assessment, this can add a crucial dimension to our understanding of neurological manifestations. While normative models serve as valuable benchmarks for evaluating brain activity, extreme deviations observed in certain individuals prompt an inquiry into the potential clinical significance of these aberrations. Such deviations may signify unique neural signatures associated with severe symptomatology or treatment-resistant cases within psychiatric disorders. The identification and examination of these extreme EEG patterns may also offer an opportunity to delineate subgroups within diagnostic categories, potentially informing personalized therapeutic interventions. However, it is imperative to approach these findings with caution, recognizing that extreme deviations may also result from individual variability, comorbidities, or methodological considerations and uncontrolled factors. Future research should delve into the nuanced clinical implications of extreme EEG deviations, striving to bridge the gap between normative modeling and real-world psychiatric presentations for a more comprehensive understanding of neurobiological substrates.

### Limitations

While our sample size is considered significantly large for EEG studies (n∼2200), it does not reach the scale often seen in MRI and fMRI studies, nor is it representative of the broader population. Furthermore, the sample size of the datasets was not uniform. The largest dataset, HBN, comprised 1539 subjects out of the total 2234 subjects considered in this study, which could bias the results. Moreover, data for this article has been sourced from five studies (detailed in <u>Methods</u>) utilizing 128-channel EEG systems. While this somewhat controls for variability stemming from the spatial resolution of the collected EEG data, it also introduces limitations to the generalizability (over other EEG systems) of the results.

One limitation of this study is the potential influence of psychiatric medication on EEG results in the assessed psychiatric patient cohort. While our research endeavors to elucidate neurobiological patterns, it is essential to recognize that medication-induced effects may introduce confounding variables. Psychotropic medications commonly prescribed for symptom management could impact the recorded EEG signals, potentially complicating the attribution of observed changes solely to underlying psychiatric conditions. Individual variations and medication interactions remain challenging to fully account for. Future studies may benefit from more extensive participant profiling, including detailed pharmacological histories (when available), to enable a more nuanced analysis of the interplay between medication and EEG outcomes.

Another challenge of the study stems from the inherent heterogeneity that may exist within each group of patients who present multiple psychiatric diagnoses. This comorbidity introduces a layer of complexity. The presence of comorbidities poses challenges in isolating the unique contributions of each condition. This complexity is reflective of the clinical reality where patients frequently exhibit overlapping symptomatology, necessitating a comprehensive approach to diagnosis and treatment. We were aware of this key point when quantifying heterogeneity and we provided an additional control analysis by investigating heterogeneity among patients who have only one diagnosis (according to the HBN dataset). The overlap results of the functional connectivity were always low and did not exceed 30%. This confirms that the observed heterogeneity is indeed intrinsic to the disorder and not driven by the possible comorbidity. In addition, we have controlled for some available parameters such as IQ (Fig. S12-S14) and sex (Fig. S2-S11, table S1-S2). Nevertheless, other factors for which information was not available for all datasets, such as the sleep quality and time of recording, may also have an impact on brain activity and should be the subject of further research and control.

Our choice of the features to model was based on the most dominant features in the literature, however, it is plausible that alternative EEG features not included in our analysis could reveal greater homogeneity among the clinical groups. A potential further analysis is to use other EEG features or combine other modalities such as MRI with EEG. Presently, the capability to train normative models for multiple response variables using GAMLSS is unavailable, thereby limiting modeling versatility. This constraint manifests in several ways. Firstly, individual models must be trained for each channel, connection, or region, escalating the complexity of both the procedure and its transition into a clinical tool. Secondly, these models are calibrated to the average values of respective channels, connections, or regions, potentially resulting in fitting inaccuracies for each model. Lastly, and notably, the inclusion of multiple response variables could yield a more rigorous representation of the deviation at the subject level.

## Conclusion

Our investigation of the electrophysiological heterogeneity across 4 psychiatric disorders reveals that EEG spectral and connectivity deviations from a normative population are extremely heterogeneous. Our findings emphasize the urgency of going *beyond the average brain* and adopting innovative EEG (and more broadly neuroimaging) approaches at the patient level, steering the field toward precision psychiatry. The complex tapestry of individual differences in EEG signatures underscores the inadequacy of current *one-size-fits-all* approaches. The call for tailored, patient-specific interventions becomes more pronounced as we navigate the intricate terrain of psychiatric heterogeneity, ultimately striving for a paradigm shift in the way we approach and understand these complex disorders.

## Methods

### Dataset

Our cohort consisted of 2234 individuals, subdivided into a group of healthy controls (n=448 in the training set, n=112 in the held-out testing set) and a group of 1674 participants clinically diagnosed with psychiatric disorders, including ADHD (n=650), ASD (n=576), ANX (n=216), and LD (n=232). The data were aggregated from five distinct studies: the Healthy Brain Network Dataset (*HBN*) (Alexander et al. 2017; Langer et al. 2017) (https://fcon_1000.projects.nitrc.org/indi/cmi_healthy_brain_network/index.html), Multimodal Resource for Studying Information Processing in the Developing Brain (*MIPDB*) (https://fcon_1000.projects.nitrc.org/indi/cmi_eeg/index.html) (Langer et al. 2017), Autism Biomarker Consortium for Clinical Trials Dataset (*ABCCT*) (https://nda.nih.gov/edit_collection.html?id=2288) (McPartland et al. 2020), Multimodal Developmental Neurogenetics of Females with ASD (*femaleASD*) (https://nda.nih.gov/edit_collection.html?id=2021) (Pelphrey 2014), and *LausanneASD*. Subjects included in this study were aged between 5 and 18 years old (mean = 9.99 ± 3.06; 45% M). High-density (128-channels) resting-state EEG data were recorded while participants had their eyes open. For a comprehensive overview of the datasets, please refer to the Supplementary Materials.

### Data Preprocessing

The EEG preprocessing and artifact removal pipeline is executed through a multi-stage, fully automated algorithm. Initially, EEG signals undergo bandpass filtering between 1 and 100 Hz, focusing on the relevant frequency range for subsequent analysis. Signals were downsampled to 200 Hz. Bad EEG channels are identified using the *pyprep* algorithm, which employs a *RANSAC*-based approach, and these channels are subsequently interpolated using information from neighboring electrodes (Bigdely-Shamlo et al. 2015; Appelhoff et al. 2022). *RANSAC* works by randomly selecting a small group of EEG channels, estimating a model based on these channels, and then identifying channels that deviate from the model as potential outliers or bad channels. This process is repeated to find a model that best fits the majority of channels while disregarding outliers. Then, re-referencing is performed using the common average reference method to minimize common noise across electrodes. Independent Component Analysis (ICA) is then applied, and the *IClabel* algorithm automatically identifies and rejects components related to eye blinks (Pion-Tonachini, Kreutz-Delgado, and Makeig 2019). A second bandpass filter narrows the frequency range to 1-45 Hz, refining the data further. The EEG signals are segmented into 10-second epochs based on experimental paradigms (e.g., eyes-open and eyes-closed). The *Autoreject* toolbox (Jas et al. 2017) is utilized for the detection and cleaning/rejection of bad epochs, ensuring the removal of artifacts or irregularities. All EEG datasets underwent the preprocessing steps described above, except when certain procedures were deemed infeasible. Notably, the Autoreject step was excluded from the preprocessing of the *ABCCT* dataset. The *femaleASD* dataset had already been preprocessed and segmented into 2-second epochs. Therefore, further preprocessing for this dataset was confined solely to downsampling and re-referencing.

### Features Extraction

#### Spectral features

As previously stated, a normative model estimates the relationship between a response variable and one or more covariates. In the context of this study, we are interested in the spectral features of the EEG signal as the designated response variable. This choice was motivated by the large literature about the alterations of EEG power in psychiatric disorders (Newson and Thiagarajan 2019). The power spectrum density (PSD) for each epoch and each channel is computed using Welch’s method (1-second *Hann* window with a 50% overlap, and a spectral resolution of 0.5 Hz). PSDs are averaged across all epochs within a single subject. To assess the relative power in specific frequency bands (delta [1-4 Hz], theta [4-8 Hz], alpha [8-13 Hz], beta [13-30 Hz], gamma [30-45 Hz]), the absolute power within each narrow band is divided by the power within the broader band [1-45 Hz].

#### Functional connectivity

EEG-based functional networks were computed using the HD-EEG source connectivity method, as described in (Hassan and Wendling 2018). Briefly, cortical sources are computed using the exact low-resolution brain electromagnetic tomography (eLORETA) which aims to reconstruct the cortical activity from EEG data with correct localization (Pascual-Marqui 2007). In our case, the noise covariance matrix was set to the identity matrix, and the regularization parameter λ was set to 0.1 (inversely proportional to the signal-to-noise ratio). Age-specific head models of the three layers (brain, skull, and scalp) were built using an MRI template of young children (4-18y) (Fonov et al. 2011). We used the Boundary Element Method (BEM) provided by the MNE Python package. The forward and inverse models were solved within a source space of 4098 sources per hemisphere (with a ∼5mm spacing between sources). Then, to streamline the complexity of the cortical sources, we downsampled them to 68 representative sources by averaging the sources within each region as defined by the Desikan-Killiany atlas (Desikan et al. 2006). Subsequently, we computed functional connectivity between pairwise regions of interest, using the amplitude envelope correlation (AEC) method, defined as the Pearson correlation between signals’ envelopes derived from the Hilbert transform (Brookes et al. 2011; Hipp et al. 2012). Zero-lag signal overlaps caused by spatial leakage were removed using a pairwise orthogonalization approach before connectivity computation (Brookes, Woolrich, and Barnes 2012).

The total number of subjects who completed the preprocessing and feature extraction steps comprises 624 individuals from the healthy control (HC) group and 2478 from the clinical group, which includes 604 subjects with ASD, 1314 with ADHD, 323 with anxiety, and 237 with learning disorders. Only the subjects between 5 and 18 years old were used in the training and testing phases. The clinical groups, namely ADHD, anxiety, and learning, were downsampled while ensuring a balanced representation across age, sex, and site/study covariates. This resulted in the final dataset used for testing, as described in the <u>data description</u> section.

### Normative Modeling

Normative Modeling (NM) seeks to establish a standard or normative relationship between a response variable (behavioral, demographic, or clinical variables) and at least one covariate (a quantitative biological measure, e.g. age, sex). In this context, Generalized Additive Models for Location, Scale, and Shape (GAMLSS) (Rigby and Stasinopoulos 2005), are semi-parametric regression models. In these models, the response variable is presumed to follow a specific distribution, wherein the parameters of this distribution can be linked to a set of explanatory variables via linear or nonlinear predictor functions, providing a flexible framework to capture complex relationships. The mathematical formulation of GAMLSS is as follows:

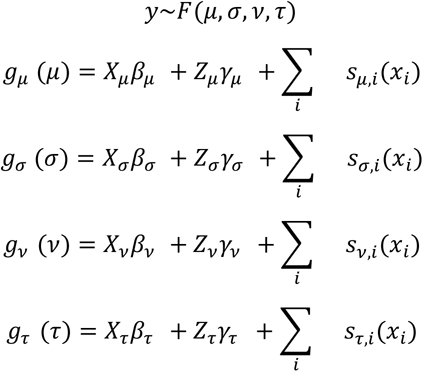

The response variable *y* is assumed to follow a distribution *F* characterized by the parameters (*μ*, *σ*, *ν*, *τ*). Each parameter can be linked to explanatory variables through the link function *g*(), where *β* represents the fixed effect term and *X* is its design matrix. *γ* accounts for the random effects, and Z is its design matrix. *s* is the non-parametric smoothing function (Rigby and Stasinopoulos 2005; Bethlehem et al. 2022). In this study, our response variable is an EEG-derived feature and the main covariate is age. The possibility of adding other covariates such as sex and data collection sites is detailed in the following sections. Bethlehem et al. use fractional polynomials as the smoothing function to account for nonlinearity without adding instability to their models (Bethlehem et al. 2022), and we adopt the same approach.

### Model distribution

GAMLSS framework offers a comprehensive list of distribution families. An empirical methodology was utilized to determine the most suitable distribution. The selection process involved training models across all considered distribution families (number of moments = 3 or more, continuous/mixed), with the Bayesian Information Criterion (BIC) serving as the comparative metric. The optimal distribution was identified as the one yielding the lowest BIC score. This selection process was systematically applied to the two features considered in this study. Distributions yielding the best fit of the averaged spectral power and averaged connectivity values are reported in Table S3 in the Supplementary Materials, respectively. The ideal number of polynomials for the age covariate and whether to consider its inclusion in parameters beyond μ is also determined based on comparing BIC scores across various models.

### Model covariates

The selection of model covariates beyond age (sex, and site/study as both a fixed effect and a random effect) is performed empirically. Each covariate is sequentially integrated into the parameter formulas. Next, the models are compared based on their BIC scores. The model yielding the lowest BIC score is selected, determining whether the covariates are retained in the final model. Subsequently, testing for the optimal distribution as described in the previous section is conducted again to reassess the suitability of the chosen distribution families. The final models for spectral and connectivity features are reported in Table S4-S5 in the Supplementary Materials.

### Model performance

#### Residual Plots

To assess the performance of our models, as recommended by XX we inspected their Q-Q plots. A good model is characterized by the presence of randomly scattered residuals around the horizontal zero line. Additionally, the kernel density estimate of the residuals should approximately follow a normal distribution, and an ideal Q-Q plot should exhibit linearity (Stasinopoulos et al. 2017). Upon reviewing our models, it appears that they demonstrate satisfactory fit and quality, Fig S15-S16 in Supplementary Materials.

#### Bootstrap analysis

The robustness and reliability of our model were assessed using bootstrap analysis. The original training set was resampled 1000 times with replacement, with the model being retrained on each of these resampled datasets. A distribution profile for the predicted median was constructed, capturing the inherent variability and stability of our model’s performance. To offer a quantifiable measure of reliability, the 95% confidence intervals were calculated from the distribution profiles, Fig. S17-S18 in Supplementary Materials. These intervals provide a statistical boundary within which the true parameter values are likely to lie, offering a clear, concise depiction of the model’s precision and stability across multiple resampling iterations.

### Deviation scores / Overlap maps

After selecting and validating the model parameters, we trained GAMLSS models for each channel/connection across all frequency bands for the reference healthy group. Subsequently, we project the test data (i.e., relative power and functional connectivity values), of our clinical groups (ADHD, ASD, ANX, LD) and the held-out healthy control group HC(test), onto the corresponding models. This process enables us to calculate deviation scores, known as z-scores (normalized quantile residuals, (Dunn and Smyth 1996)), for each channel/connection for each subject, resulting in an individual deviation map for each subject. An extreme deviation is defined as |z-score| > 2. Consequently, we derived positive and negative extreme deviation maps for z-scores > 2 and < -2, respectively. We then computed the number of subjects exhibiting at least one extreme deviation, as well as the number of extreme deviations per subject. Additionally, for each channel/connection, we assessed the percentage of subjects exhibiting extreme deviation at that location among those with at least one extreme deviation, leading to an overlap map of the extreme deviation in each group.

### Permutation test

We used group-based permutation tests to evaluate group differences in channel/connectivity-level overlap (Segal et al. 2023). These tests involved shuffling cases and control labels of the individual-specific deviation maps. At each iteration, we permuted group labels and obtained a new grouping of extreme deviation maps for each subject based on the shuffled labels. Subsequently, new overlap maps were computed for HC(test) and each clinical group. We then subtracted the surrogate HC(test) overlap map from the surrogate clinical group’s overlap map to derive an overlap difference map for each disorder. This procedure was repeated 5,000 times to establish an empirical distribution of overlap difference maps under the null hypothesis of random group assignment. Finally, for each channel/connection, we obtained *p*-values as the proportion of null values that exceeded the observed difference. Statistically significant effects were identified using two-tailed FDR correction (*p<0.05)*.

### Data availability

The Healthy Brain Network (*HBN*) dataset is available at https://fcon_1000.projects.nitrc.org/indi/cmi_healthy_brain_network/index.html. The Multimodal Resource for Studying Information Processing in the Developing Brain (*MIPDB*) Dataset is accessible at https://fcon_1000.projects.nitrc.org/indi/cmi_eeg/index.html. The Autism Biomarker Consortium for Clinical Trials Dataset (*ABCCT*) and the Multimodal Developmental Neurogenetics of Females with ASD (*femaleASD*) can be requested from the NIMH Data Archive platform (https://nda.nih.gov/edit_collection.html?id=2288, https://nda.nih.gov/edit_collection.html?id=2021). The *LausanneASD* dataset is available upon request from [BR].

### Code availability

Codes are available at https://github.com/MINDIG-1/NM-Psy.git. For statistical modeling, we employed the gamlss package in R, (Mikis Stasinopoulos and Rigby 2008). EEG signal processing is done using the MNE-python package (https://mne.tools/stable/index.html). Networks are visualized using BrainNet Viewer (https://www.nitrc.org/projects/bnv/) (Xia, Wang, and He 2013). Network comparisons were performed utilizing the network-based statistic (NBS) tool, (https://www.nitrc.org/projects/nbs) (Zalesky, Fornito, and Bullmore 2010).

## Acknowledgements

This work was fully funded by MINDIG as a part of its R&D activity. We would like to thank all the researchers who shared their data in open-access and all the participants (patients and controls) who approved the use of their data in research.

## Author contributions

A.E., S.A., and M.H. conceived the study and wrote the manuscript, with valuable revision from all authors. Results were interpreted by A.E., S.A., and M.H., with contributions from G.R., A.L., and A.I.. Figures were produced by A.M and A.E.. B.R. and N.C. provided ASD data. J.T. and A.K. helped in the data preprocessing and analysis part.

