## supplementary materials for "Beyond homogeneity: Charting the landscape of heterogeneity in psychiatric electroencephalography"

### Methods

#### Datasets

Our cohort consisted of 2234 individuals, subdivided into a group of healthy controls (n=448 in the training set, n=112 in the held-out testing set) and a group of 1674 participants clinically diagnosed with psychiatric disorders, including ADHD (n=650), ASD (n=576), anxiety disorder (n=216), and learning disorder (n=232). The data were aggregated from five distinct studies, each detailed subsequently.

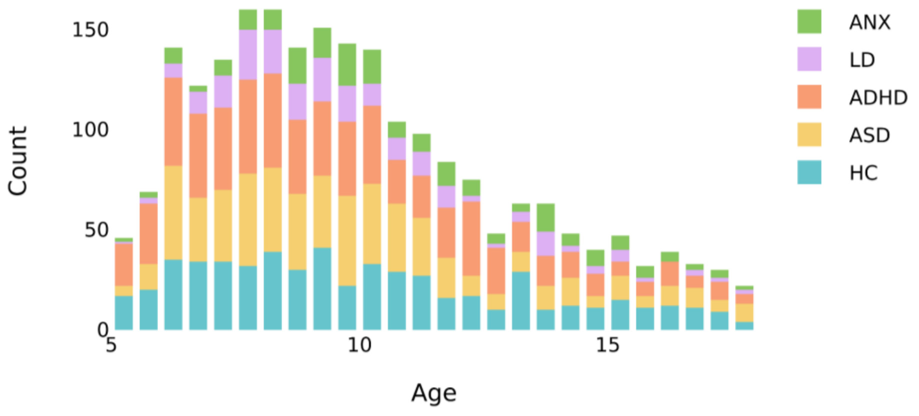


**Figure. S1 | Age distribution across groups.**

**Dataset 1: Healthy Brain Network Dataset (*HBN*)**

***Participants.*** The Healthy Brain Network (*HBN*) project (https://fcon_1000.projects.nitrc.org/indi/cmi_healthy_brain_network/index.html) (Alexander et al. 2017; Langer et al. 2017), initiated by the Child Mind Institute, enrolled children and adolescents aged 5–21 from four study sites in the New York City area. Inclusion criteria required participants to possess adequate verbal communication skills, with assistance from parents or guardians when needed. Exclusion criteria encompassed severe neurological disorders, significant cognitive impairment (IQ<66), acute encephalopathy, known neurodegenerative disorders, or other abnormalities hindering complete participation in the protocol. A comprehensive set of neuropsychological assessments was administered (for details, see Ref). The dataset encompasses 3234 participants, categorized into healthy controls and clinical groups predominantly associated with neurodevelopmental disorders, including ADHD, ASD, learning disorders, anxiety, and obsessive-compulsive disorder, among others.

***EEG Acquisition.*** High-density EEG data was recorded in an acoustically isolated room using a 128-channel EEG Geodesic HydroCel system provided by Electrical Geodesics Inc. (EGI) during a resting-state paradigm, where participants were instructed to fixate on a cross displayed at the center of a computer screen and to open or close their eyes at different time points. The recordings were conducted at a sampling rate of 500Hz, with a bandpass filter between 0.1 and 100Hz, and a recording reference at Cz (head vertex). Electrode impedance checks were performed before recording, and values were maintained below 40 kΩ. (See (Alexander et al. 2017; Langer et al. 2017) for more details).

**Dataset 2: Multimodal Resource for Studying Information Processing in the Developing Brain (*MIPDB*)**

***Participants***. The *MIPDB* dataset (https://fcon_1000.projects.nitrc.org/indi/cmi_eeg/index.html) (Langer et al. 2017) included 101 healthy controls and (n=101) and 31 patients with neurodevelopmental disorders (ADHD, ASD, learning disorders, anxiety).

***EEG Acquisition.*** Resting-state high-density EEG data were recorded at a sampling rate of 500 Hz with a bandpass of 0.1 to 100 Hz, using a 128-channel EEG Geodesic Hydrocel system. Participants were instructed to focus on a central cross and to open or close their eyes in response to an auditory beep, following a sequence of alternating 20-second blocks of eyes open and 40-second blocks of eyes closed

**Dataset 3: Autism Biomarker Consortium for Clinical Trials Dataset (*ABCCT*)**

***Participants.*** The *ABC-CT* dataset (https://nda.nih.gov/edit_collection.html?id=2288), included a total of 280 children diagnosed with Autism Spectrum Disorder (ASD) and 119 typically developing (TD) children, aged between 6 and 11 years. The diagnosis of ASD was established using the Autism Diagnostic Observation Schedule (ADOS). The Full-Scale Intelligence Quotient (FSIQ) of the subjects ranged from 60 to 150, as assessed by the Differential Ability Scales. Exclusion criteria encompassed known genetic syndromes, neurological conditions causally linked to ASD, recognized metabolic disorders, and mitochondrial dysfunction. Medications were permitted to ensure the sample's representativeness for individuals undergoing treatment, promoting the generalizability of biomarker measurements. However, a stable medication regimen (maintained for at least 8 weeks prior to enrollment) was a prerequisite for inclusion in the study. (See (McPartland et al. 2020) for more details).

***EEG Acquisition***. Eyes open resting-state EEG was recorded for 5 min during the viewing of abstract videos. The resting-state EEG data were captured using a 128-channel EEG geodesic hydrocel system manufactured by Electrical Geodesics Inc. (EGI), with recordings conducted at a sampling rate of 1000Hz. The recording reference point was set at Cz (head vertex), and electrode impedance was assessed before each recording, ensuring it remained below 100 kΩ.

**Dataset 4: Multimodal Developmental Neurogenetics of Females with ASD (*femaleASD*)**

***Participants.*** A sex-balanced cohort of ASD, matched typically developing participants, and unaffected siblings, all aged between 8 and 17 years were recruited as part of the Autism Center for Excellence (ACE) project Multimodal Developmental Neurogenetics of Females with ASD (Pelphrey 2014), focusing on sex differences in children with ASD (https://nda.nih.gov/edit_collection.html?id=2021). Of the enrolled children, 339 participated in the EEG protocol.

***EEG Acquisition.*** Resting-state EEG data were recorded using a 128-channel EGI system (Electrical Geodesics Inc.) with a 500 Hz sampling rate. The experiment session consisted of three runs, each of 2 blocks of 1 minute of eyes open to screen saver movie and 30 seconds of eyes closed.

**Dataset 5: *LausanneASD***

***Participants.*** This study enlisted a sample of 139 children (75 males and 64 females), comprising 58 participants diagnosed with ASD and 81 typically developing (TD) participants. All participants were aged between 2 and 10 years. TD children were recruited from the general population via an online form. Inclusion criteria were the following: being enrolled in the regular school system and having no known NDD or learning disabilities. Children with ASD were all recruited in the Service des Troubles du Spectre de l'Autisme et apparentés at Lausanne University Hospital, Switzerland (STSA-a). Trained psychologists and child psychiatrists established the diagnosis of ASD according to the Diagnostic and Statistical Manual of Mental Disorders, fifth edition (American Psychiatric Association, 2013). This included a review of patients' medical and developmental history, an assessment with the ADI-R, and the Autism Diagnosis Observation Scale-2 (ADOS-2; (Lord et al. 2012). In both cohorts, parents or legal guardians responding to caregiver-report questionnaires were required to be native or fluent French speakers. The study was reviewed and approved by the local Ethics committee and signed consent forms were obtained from participants or legal representatives.

***EEG Acquisition.*** EEG signals were recorded using a 128-channel EEG system (Geodesic Sensor Net -128 electrodes). The impedance of each electrode is checked prior to recording to ensure good contact and is kept below 10 kΩ. Signals were sampled at 1000 or 250 Hz. All subjects underwent a 5-minute resting-state EEG session in which they were asked to relax and keep their eyes open.

### Exploratory Data Analysis

We performed the Wilcoxon Rank-Sum test, also known as the Mann-Whitney U test, to check for any significant differences in the EEG spectral power/functional connectivity distribution between male and female participants within each diagnostic group. The effect sizes of these differences were quantified using Cohen's d.

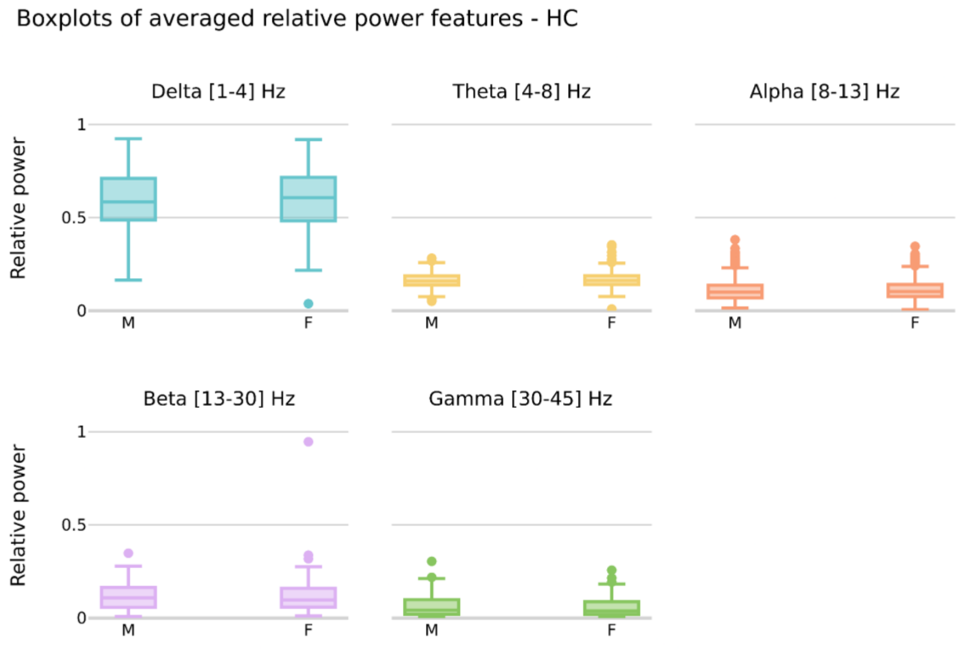


**Figure. S2 | Sex differences in averaged relative power of EEG frequency bands in HC group.** (*) denote statistically significant differences between sexes within each frequency band.

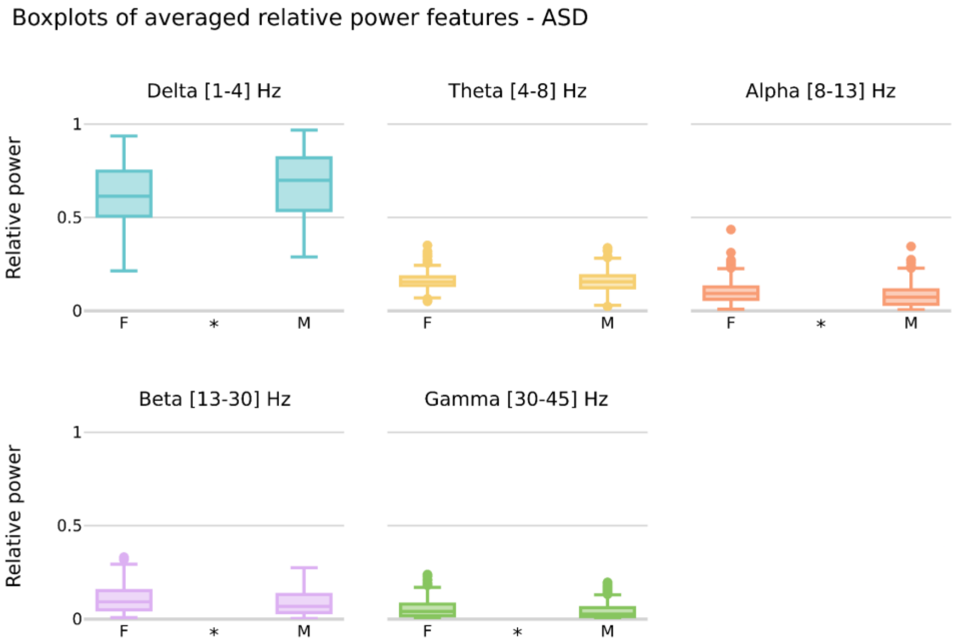


**Figure. S3 | Sex differences in averaged relative power of EEG frequency bands in ASD group.** (*) denote statistically significant differences between sexes within each frequency band.

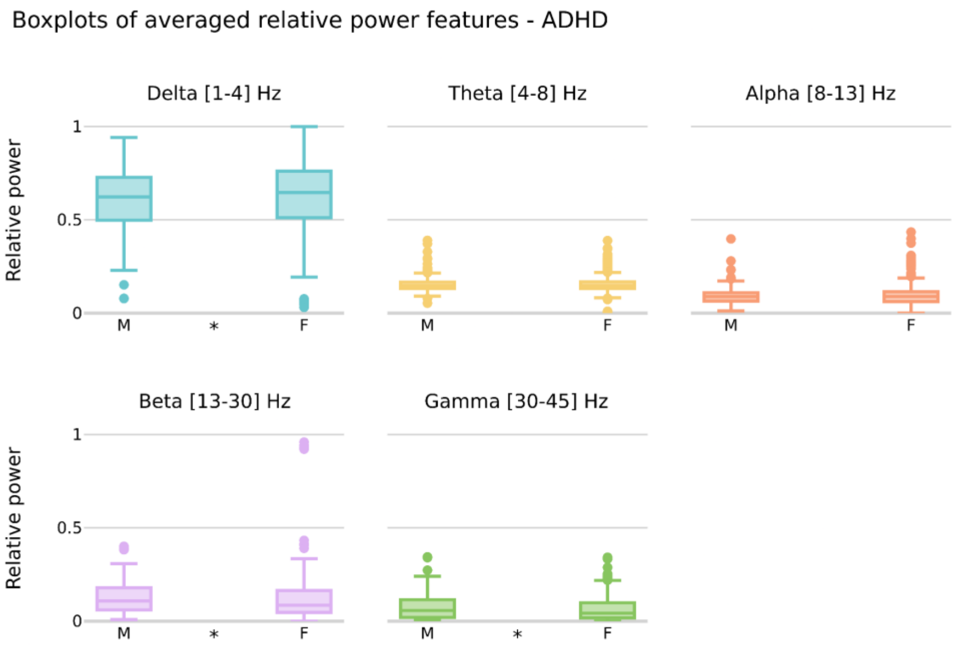

**Figure. S4 | Sex differences in averaged relative power of EEG frequency bands in ADHD group.** (*) denote statistically significant differences between sexes within each frequency band.

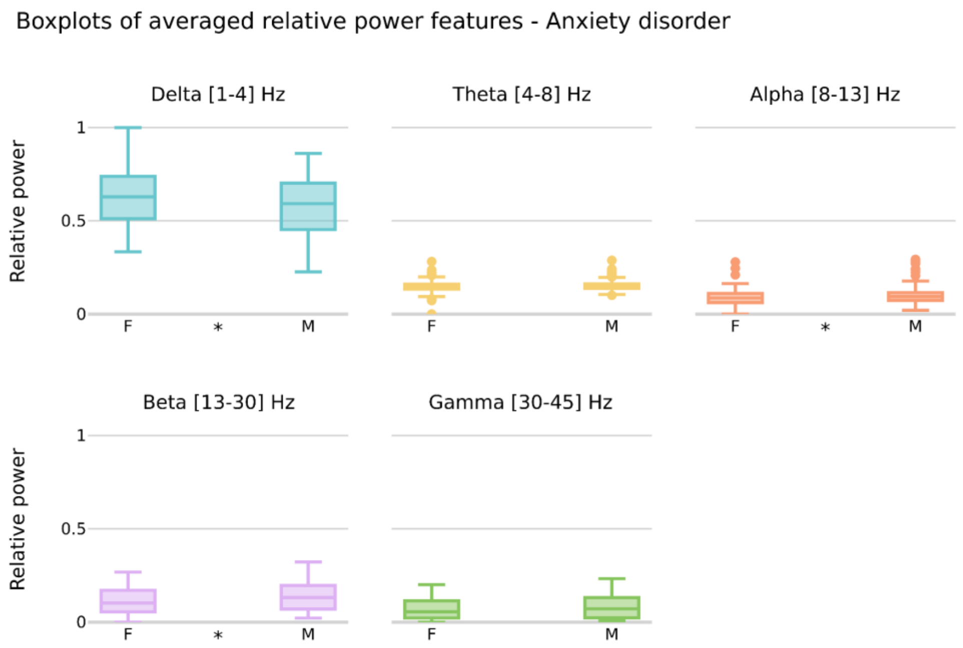


**Figure. S5 | Sex differences in averaged relative power of EEG frequency bands in ANX group.** (*) denote statistically significant differences between sexes within each frequency band.

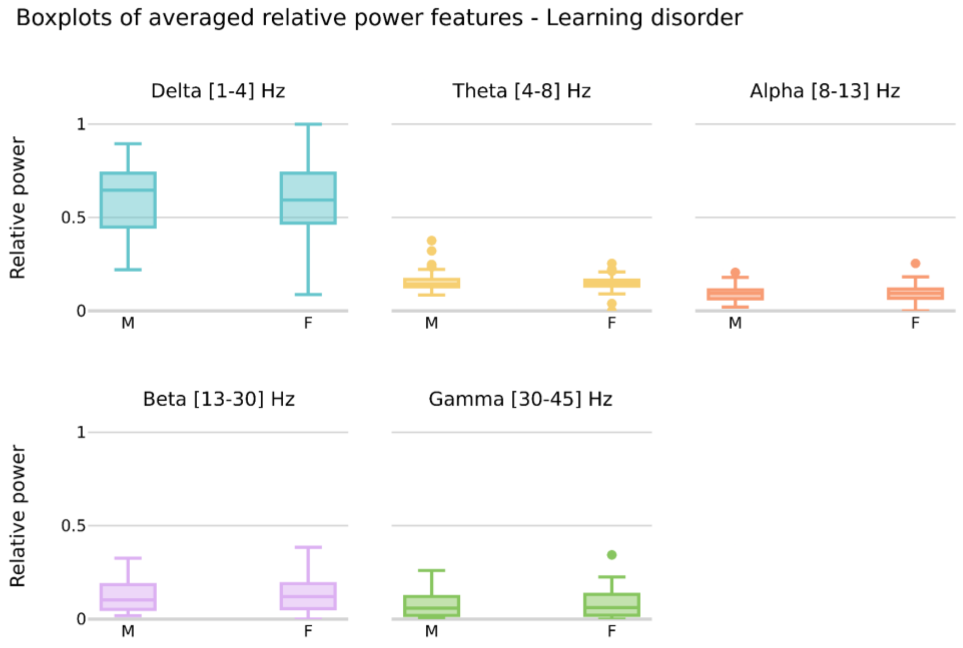


**Figure. S6 | Sex differences in averaged relative power of EEG frequency bands in LD group.** (*) denote statistically significant differences between sexes within each frequency band.

**Table. S1 | Sex differences in averaged realtive relative power of EEG frequency bands: *P-value (Cohen's d).***

| | Delta [1-4] Hz | Theta [4-8] Hz | Alpha [8-13] Hz | Beta [13-30] Hz | Gamma [30-45] Hz |
|---|---|---|---|---|---|
| **HC** | 0.48 (-0.05) | 0.50 (-0.07) | 0.41 (-0.02) | 0.27 (0.05) | 0.18 (0.14) |
| **ASD** | 0.00 (0.35) | 0.55 (-0.09) | 0.00 (-0.33) | 0.00 (-0.29) | 0.00 (-0.32) |
| **ADHD** | 0.01 (-0.13) | 0.57 (0.08) | 0.94 (-0.03) | 0.00 (0.12) | 0.01 (0.16) |
| **LD** | 0.87 (0.01) | 0.81 (0.12) | 0.96 (-0.00) | 0.72 (-0.04) | 0.78 (-0.05) |
| **ANX** | 0.01 (-0.35) | 0.48 (0.17) | 0.03 (0.27) | 0.02 (0.31) | 0.08 (0.26) |

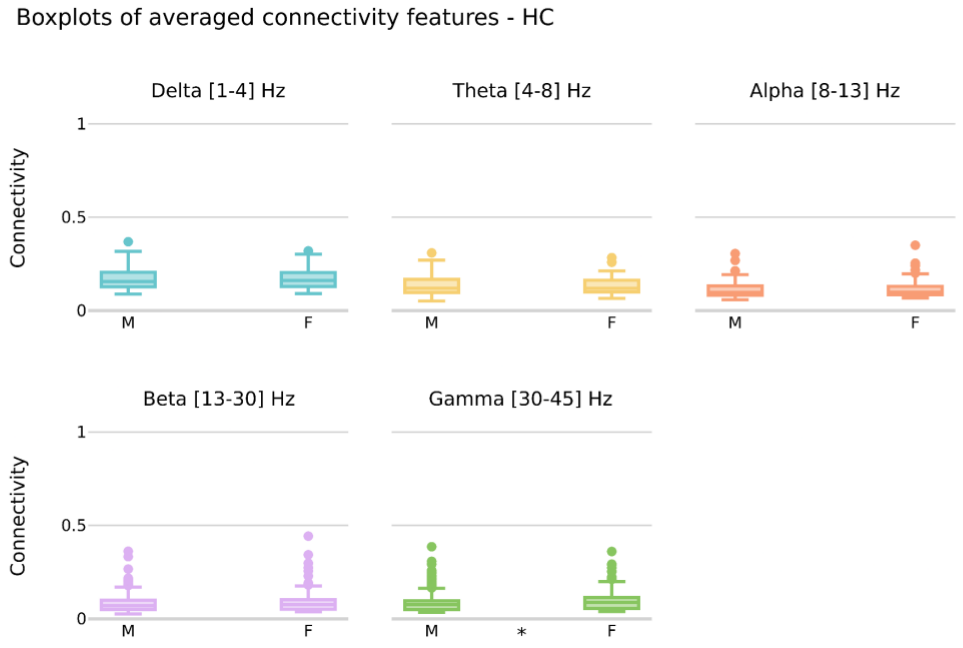


**Figure. S7 | Sex differences in averaged functional connectivity in HC group.** (*) denote statistically significant differences between sexes within each frequency band.

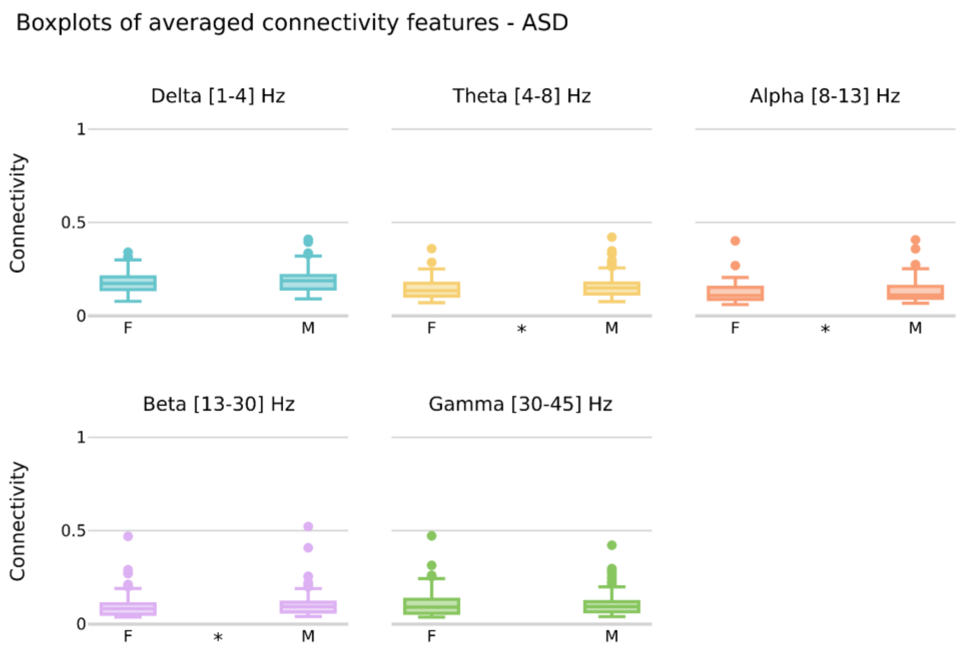


**Figure. S8 | Sex differences in averaged functional connectivity in ASD group.** (*) denote statistically significant differences between sexes within each frequency band.

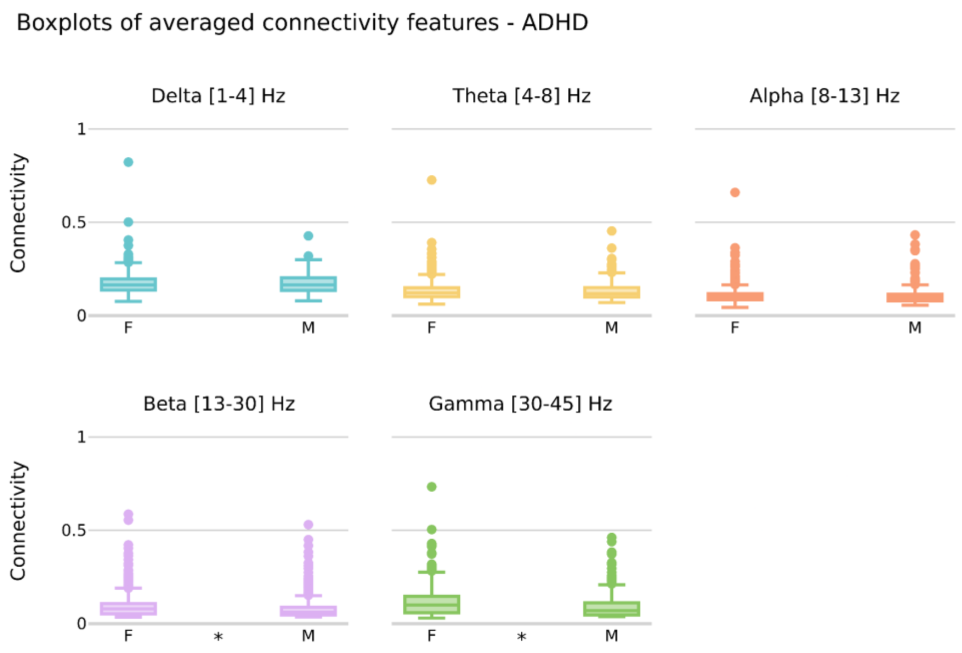

**Figure. S9 | Sex differences in averaged functional connectivity in ADHD group.** (*) denote statistically significant differences between sexes within each frequency band.

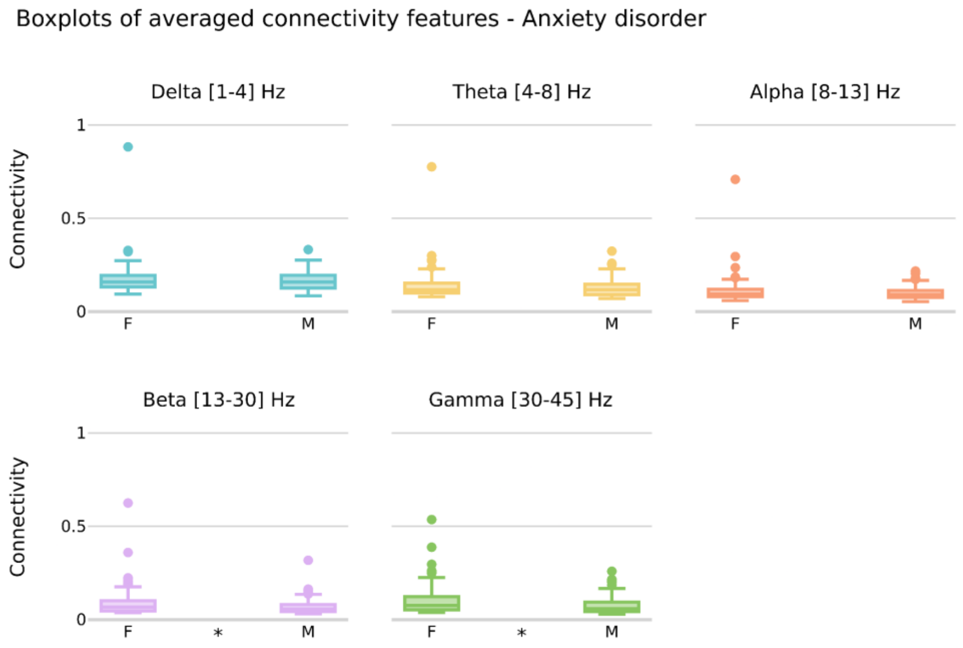


**Figure. S10 | Sex differences in averaged functional connectivity in ANX group.** (*) denote statistically significant differences between sexes within each frequency band.

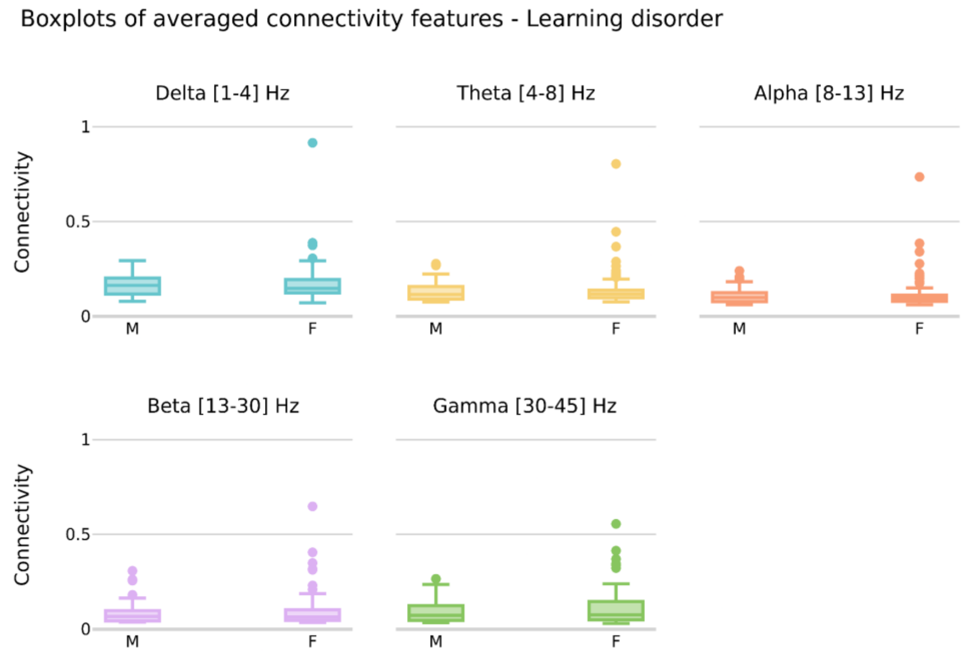


**Figure. S11 | Sex differences in averaged functional connectivity in LD group.** (*) denote statistically significant differences between sexes within each frequency band.

**Table. S2 | Sex differences in averaged relative power of EEG frequency bands:** *P-value (Cohen's d).*

| | Delta [1-4] Hz | Theta [4-8] Hz | Alpha [8-13] Hz | Beta [13-30] Hz | Gamma [30-45] Hz |
|---|---|---|---|---|---|
| HC | 0.83 (0.00) | 0.97 (0.03) | 0.62 (-0.03) | 0.27 (-0.08) | 0.00 (-0.21) |
| ADHD | 0.67 (0.03) | 0.78 (0.06) | 0.06 (0.02) | 0.00 (-0.12) | 0.00 (-0.34) |
| ASD | 0.09 (0.18) | 0.02 (0.24) | 0.05 (0.18) | 0.01 (0.15) | 0.43 (-0.01) |
| Learning | 0.50 (-0.01) | 0.80 (-0.05) | 0.66 (-0.12) | 0.54 (-0.19) | 0.29 (-0.21) |
| Anxiety | 0.55 (-0.12) | 0.39 (-0.12) | 0.19 (-0.19) | 0.00 (-0.35) | 0.00 (-0.42) |

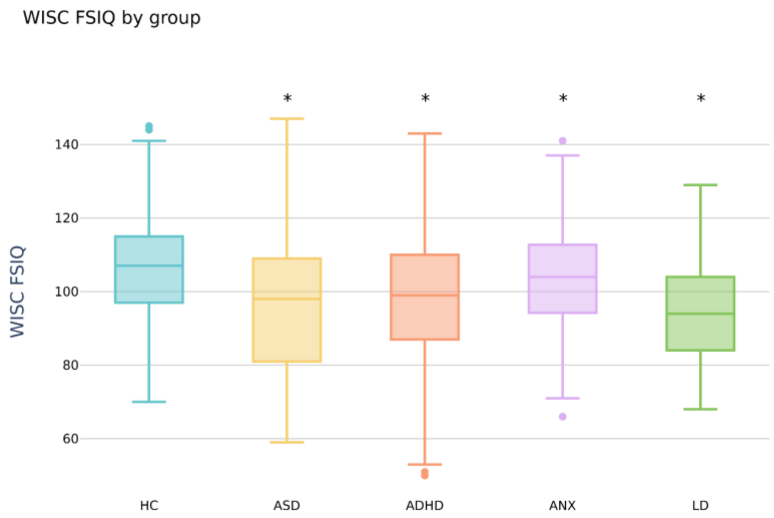


**Figure. S12 | WISQ FSIQ scores distribution across groups.** (*) denote statistically significant differences between groups.

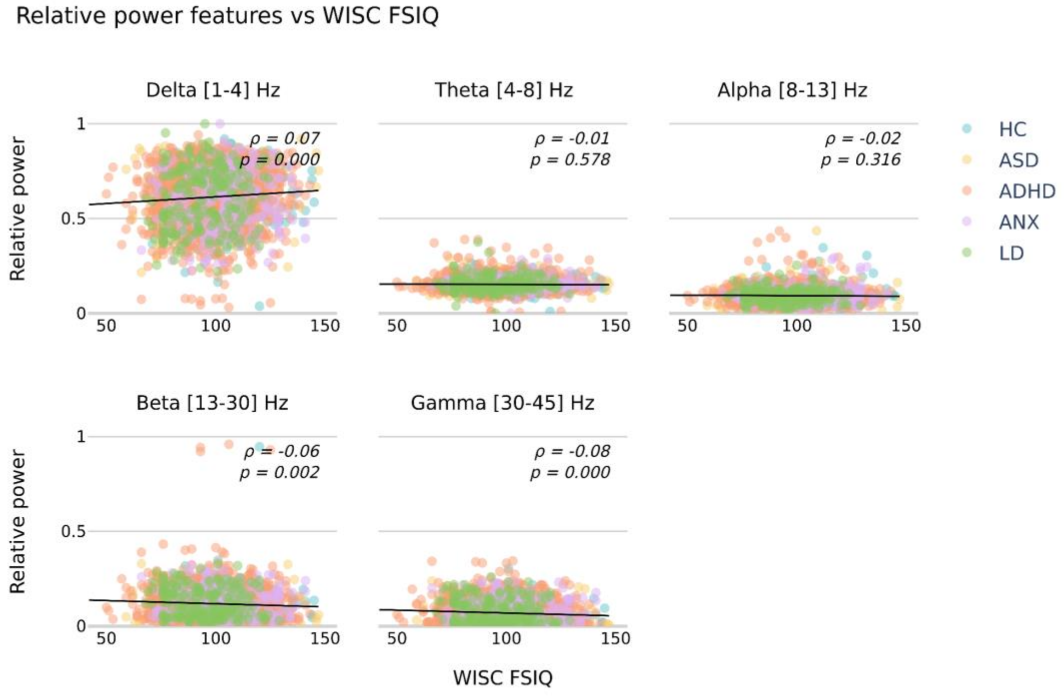


**Figure. S13 | Correlation between the average relative power and WISQ FSIQ scores**

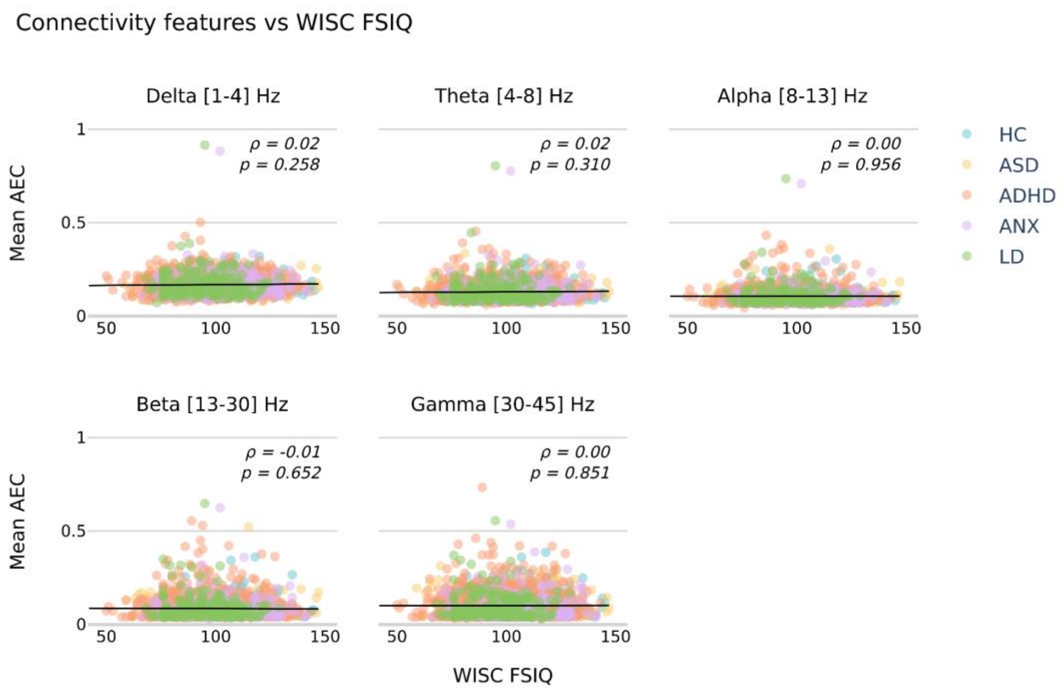


**Figure. S14 | Correlation between the average functional connectivity and WISQ FSIQ scores**

### Normative Modeling

#### Model distribution

**Table. S3 | Distribution families yielding the best fit of the averaged spectral power and averaged connectivity**

| | Frequency band | Delta | Theta | Alpha | Beta | Gamma |
|---|---|---|---|---|---|---|
| Distribution family | spectral features | GT | SEP2 | BCT | GG | exGAUS |
| | connectivity features | BCPE | BCPE | ST3 | ST3 | SEP2 |

#### Model covariates

The final models for spectral and connectivity features are reported in Table XX and Table YY.

**Table. S4 | Model equations and family distribution for spectral features models across all frequency bands**

| Frequency band | Delta | Theta | Alpha | Beta | Gamma |
|---|---|---|---|---|---|
| mu | y ~ fp (age, 1) | y ~ fp (age, 1) + factor (site) | y ~ fp (age, 1) + factor (site) | y ~ fp (age, 1) + factor (site) | y ~ fp (age, 1) |
| sigma | - | ~ factor (site) | ~ factor (site) | ~ fp (age, 1) + factor (site) | - |
| Distribution Family | SEP4 | SEP2 | GB2 | SEP1 | exGAUS |

**Table. S5 | Model equations and family distribution for connectivity features models across all frequency bands**

| Frequency band | Delta | Theta | Alpha | Beta | Gamma |
|---|---|---|---|---|---|
| mu | y ~ fp (age, 2) | y ~ fp (age, 2) | y ~ fp (age, 1) + factor (sex) | y ~ fp (age, 1) | y ~ fp (age, 1) |
| sigma | ~ fp (age, 1) | ~ fp (age, 1) | ~ fp (age, 1) | ~ fp (age, 1) | ~ fp (age, 1) |
| nu | - | ~ fp (age, 1) | - | - | - |
| Distribution Family | BCPE | BCPE | ST3 | SEP2 | SEP2 |

### Model Validation

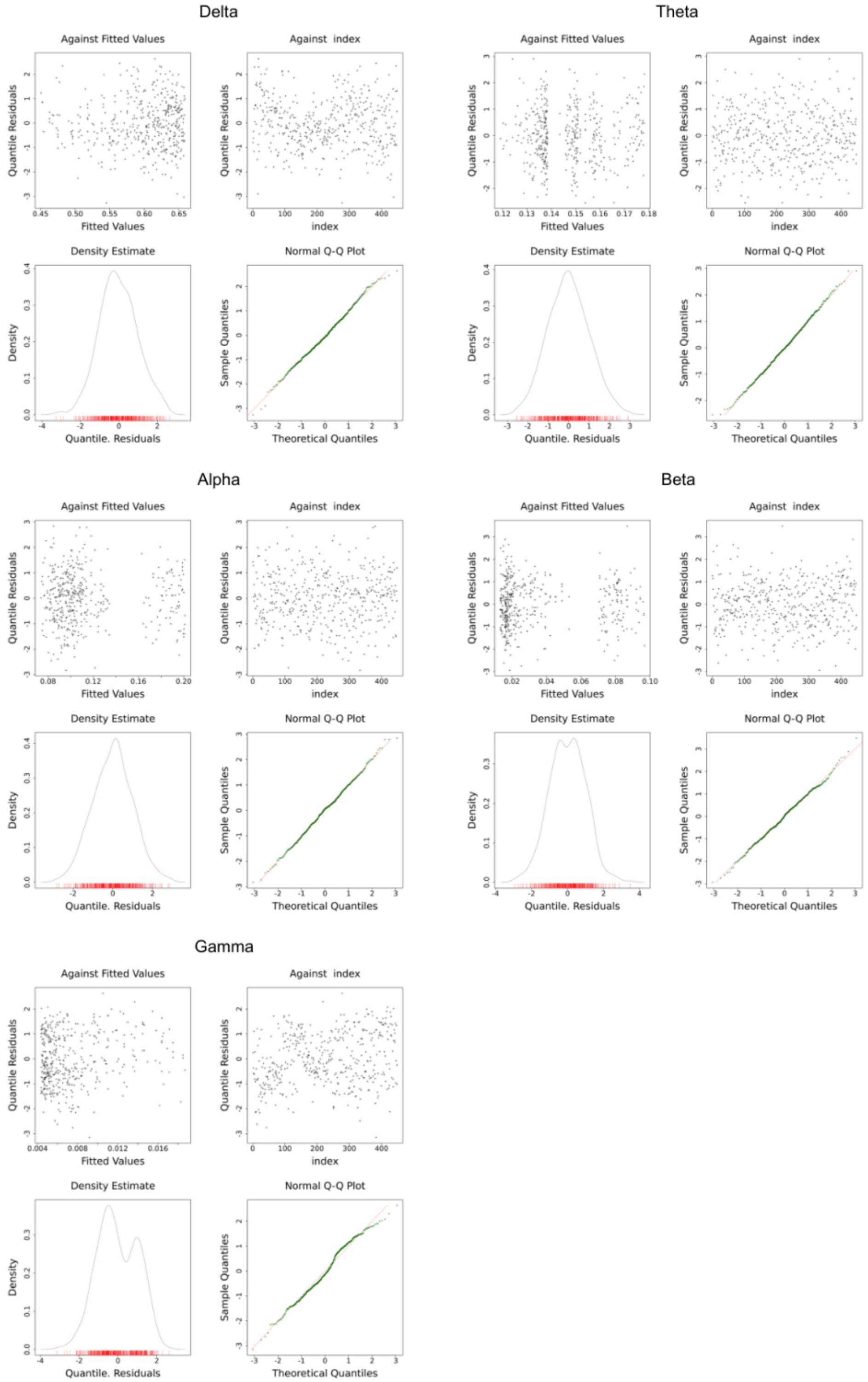


**Fig. S15 | Diagnostic residual plots for assessing spectral model fit:** residuals *vs* fitted values, index, density estimate, and normal Q-Q plot.

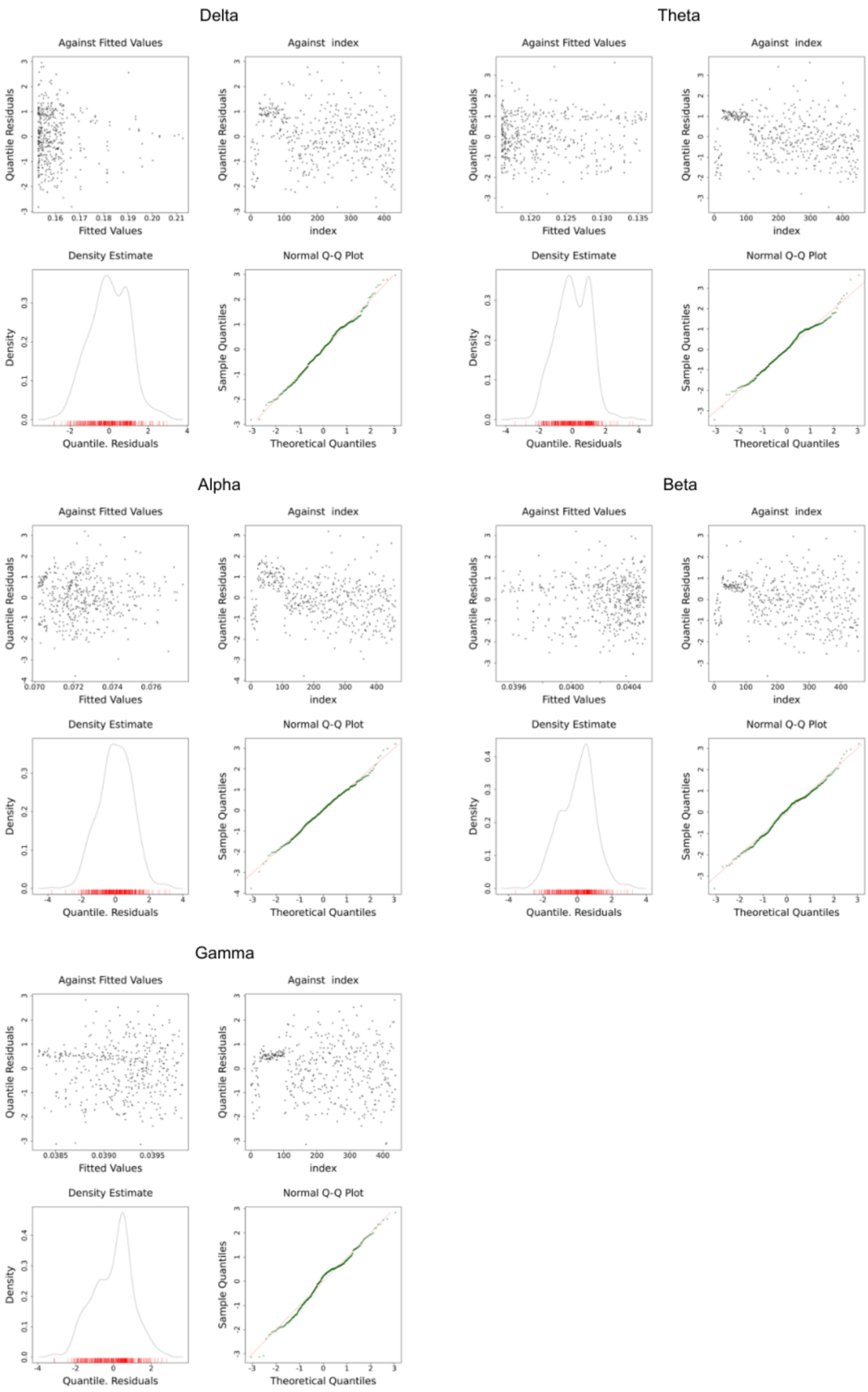


**Fig. S16 | Diagnostic residual plots for assessing connectivity model fit:** residuals *vs* fitted values, index, density estimate, and normal Q-Q plot.

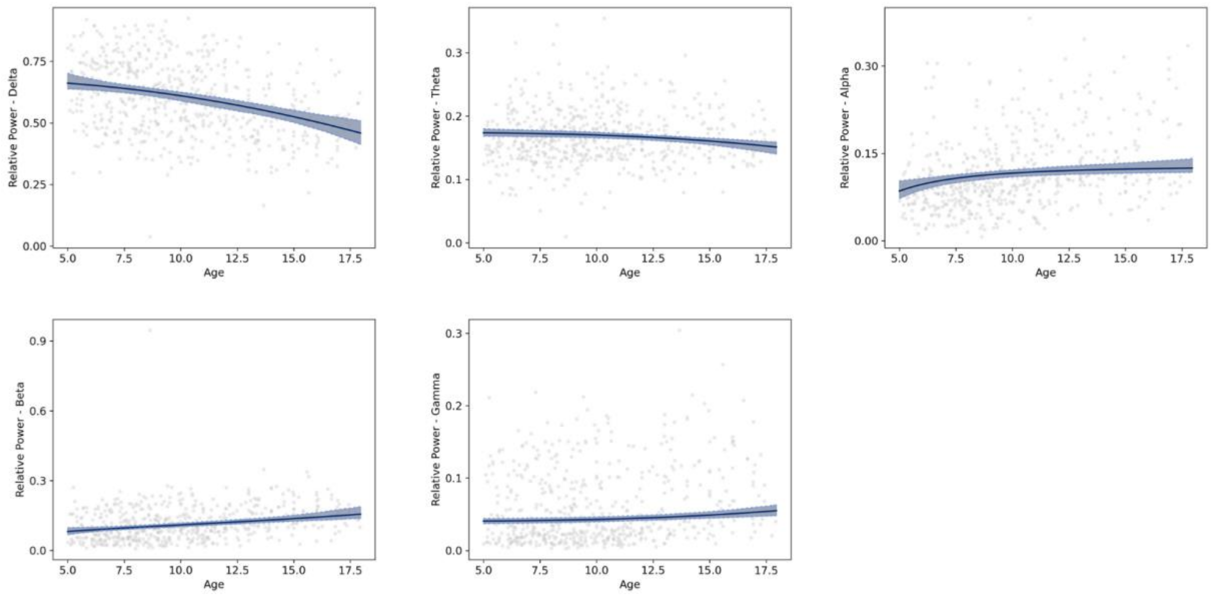


**Figure. S17 | Bootstrap analyses of spectral models.** The bold line is the model median and the shaded area represents the 95% confidence interval.

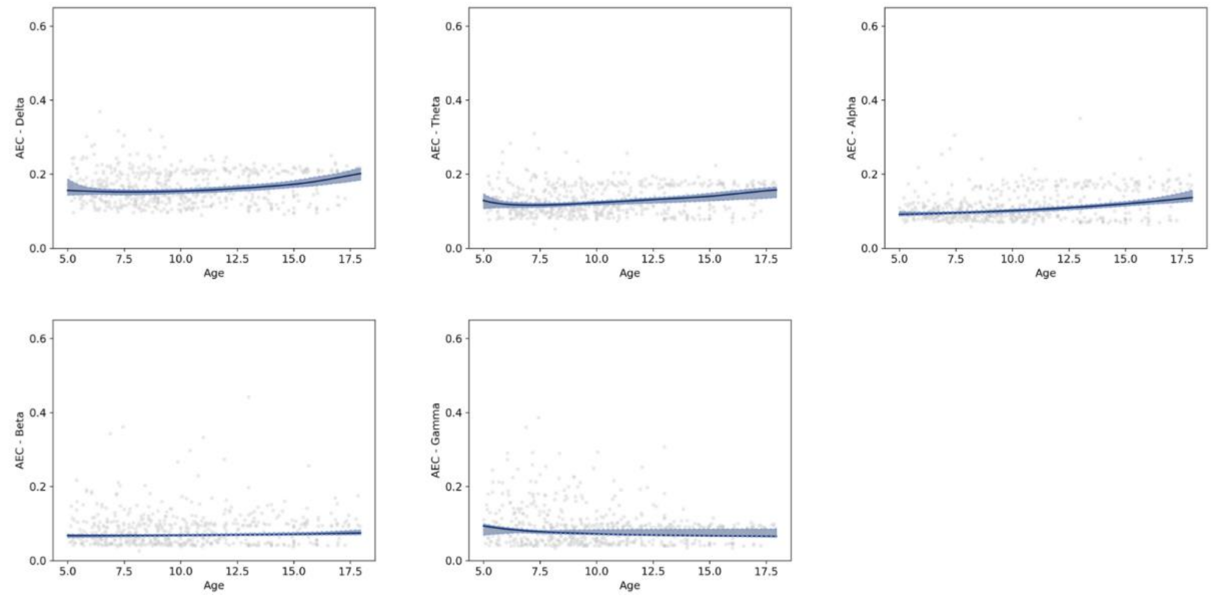


**Figure. S18 | Bootstrap analyses of connectivity models.** The bold line is the model median and the shaded area represents the 95% confidence interval.

### Results

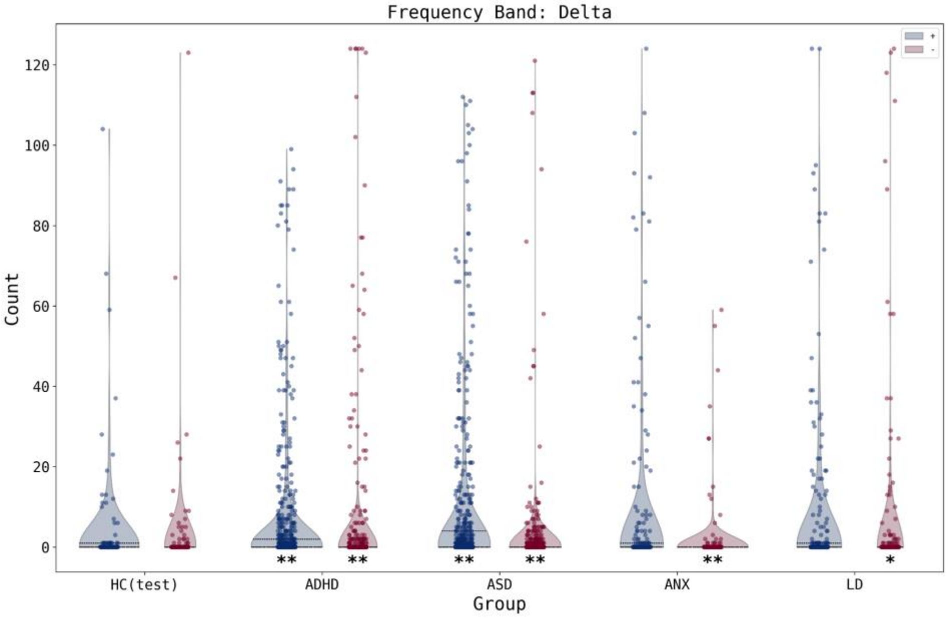


**Figure. S19 | Distribution of the number of extremely deviated channels per subject across groups in delta band.** Blue (Red) violins represent positive (negative) deviations. (*) denotes significant difference between HC and cases. (*) denotes *p<0.05*, (**) denotes *p<0.01*.

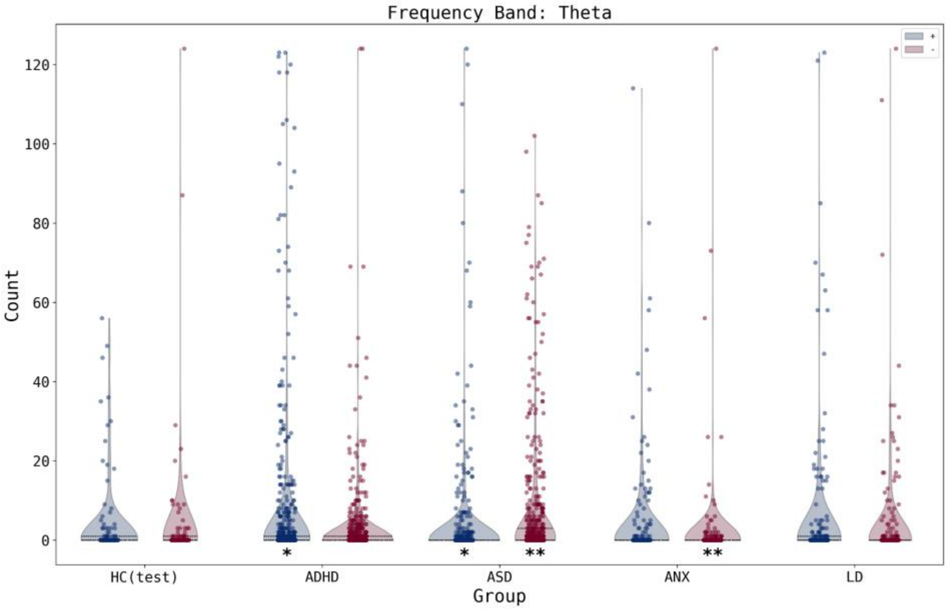


**Figure. S20 | Distribution of the number of extremely deviated channels per subject across groups in theta band.** Blue (Red) violins represent positive (negative) deviations. (*) denotes significant difference between HC and cases. (*) denotes *p<0.05*, (**) denotes *p<0.01*.

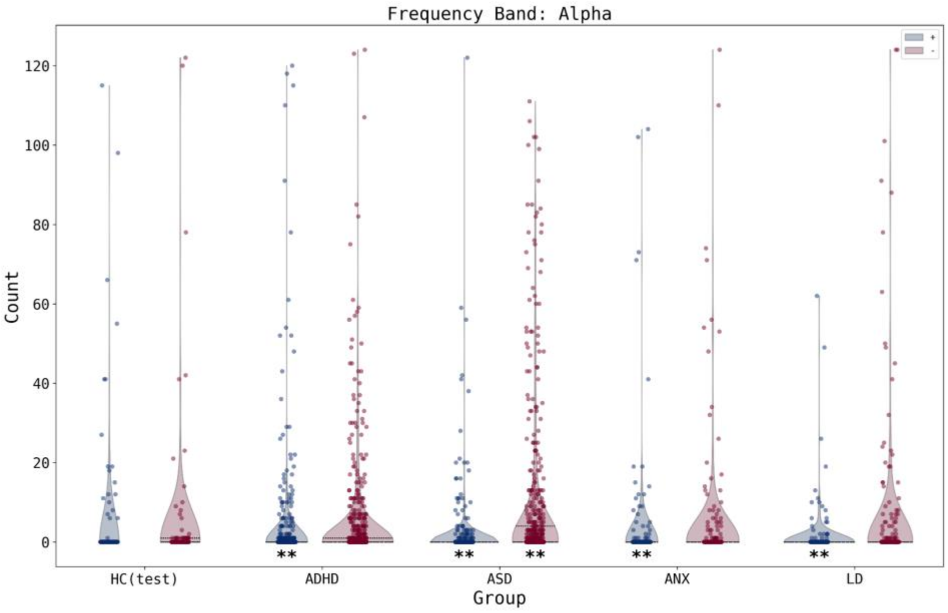


**Figure. S21 | Distribution of the number of extremely deviated channels per subject across groups in alpha band.** Blue (Red) violins represent positive (negative) deviations. (*) denotes significant difference between HC and cases. (*) denotes *p<0.05*, (**) denotes *p<0.01*.

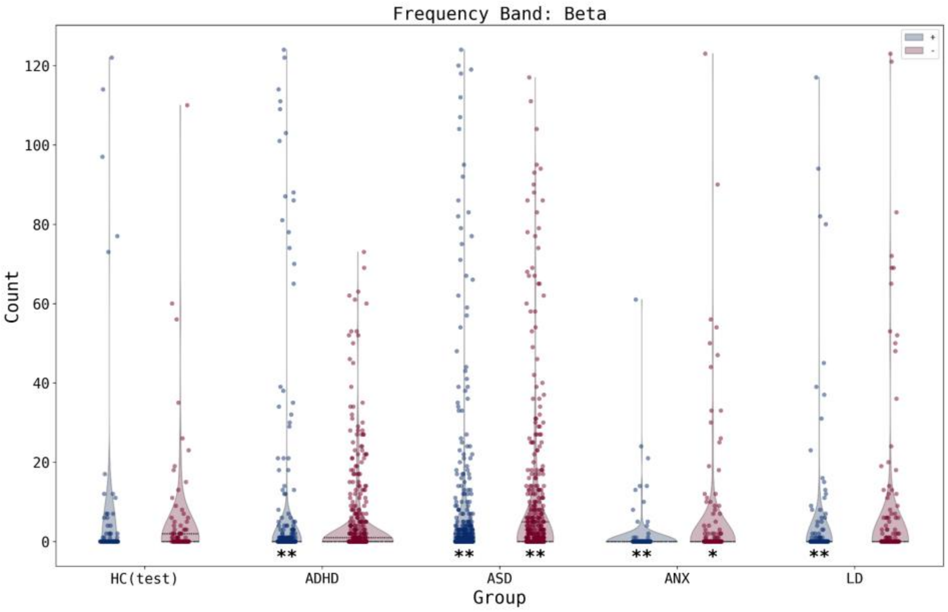


**Figure. S22 | Distribution of the number of extremely deviated channels per subject across groups in beta band.** Blue (Red) violins represent positive (negative) deviations. (*) denotes significant difference between HC and cases. (*) denotes *p<0.05*, (**) denotes *p<0.01*.

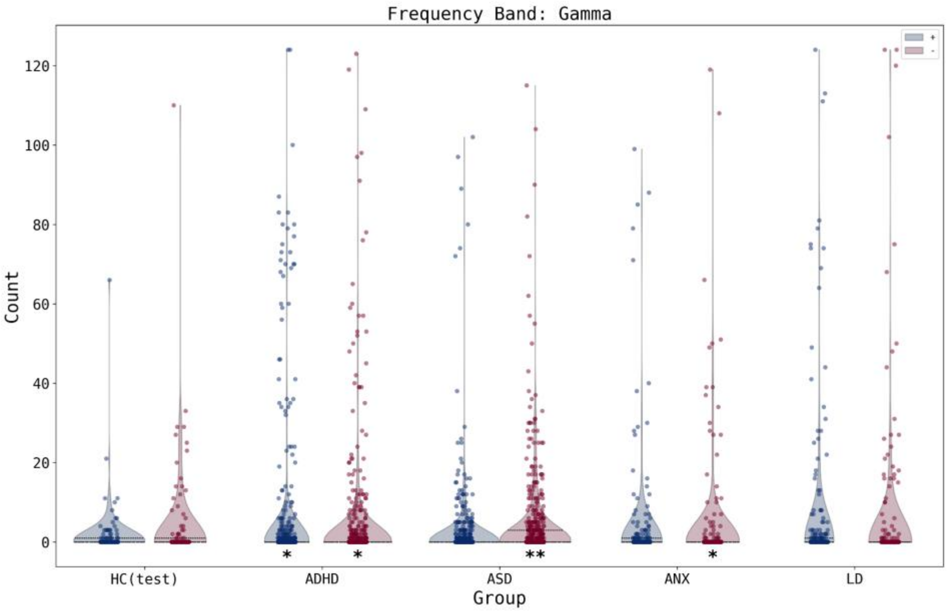


**Figure. S23 | Distribution of the number of extremely deviated channels per subject across groups in gamma band.** Blue (Red) violins represent positive (negative) deviations. (*) denotes significant difference between HC and cases. (*) denotes *p<0.05*, (**) denotes *p<0.01*.

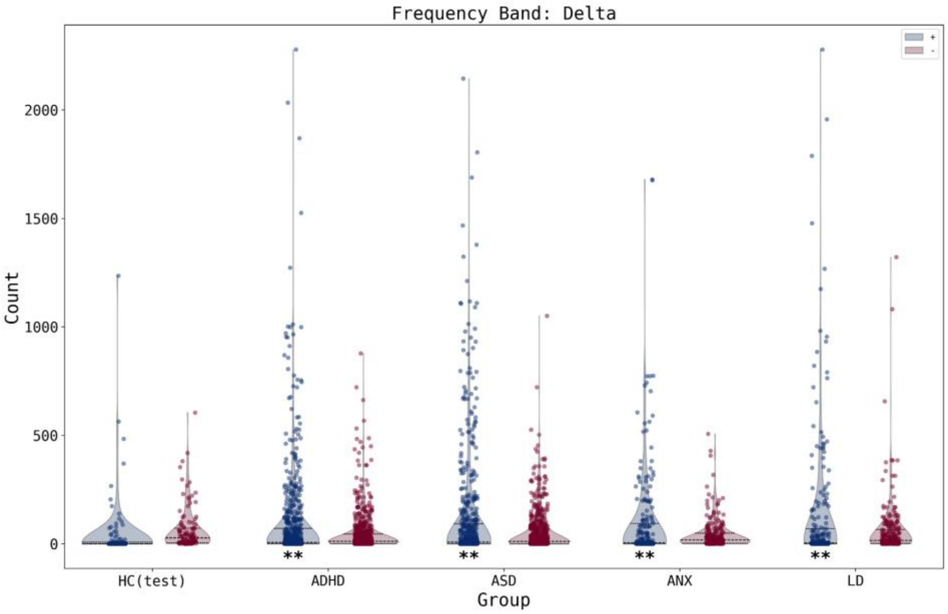


**Figure. S24 | Distribution of the number of extremely deviated connections per subject across groups in delta band.** Blue (Red) violins represent positive (negative) deviations. (*) denotes significant difference between HC and cases. (*) denotes *p<0.05*, (**) denotes *p<0.01*.

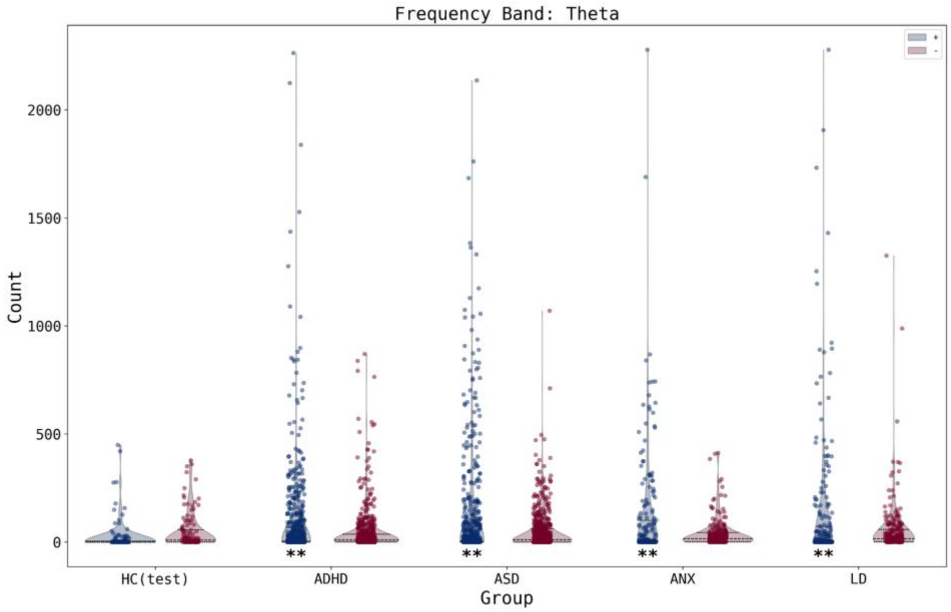


**Figure. S25 | Distribution of the number of extremely deviated connections per subject across groups in theta band.** Blue (Red) violins represent positive (negative) deviations. (*) denotes significant difference between HC and cases. (*) denotes *p<0.05*, (**) denotes *p<0.01*.

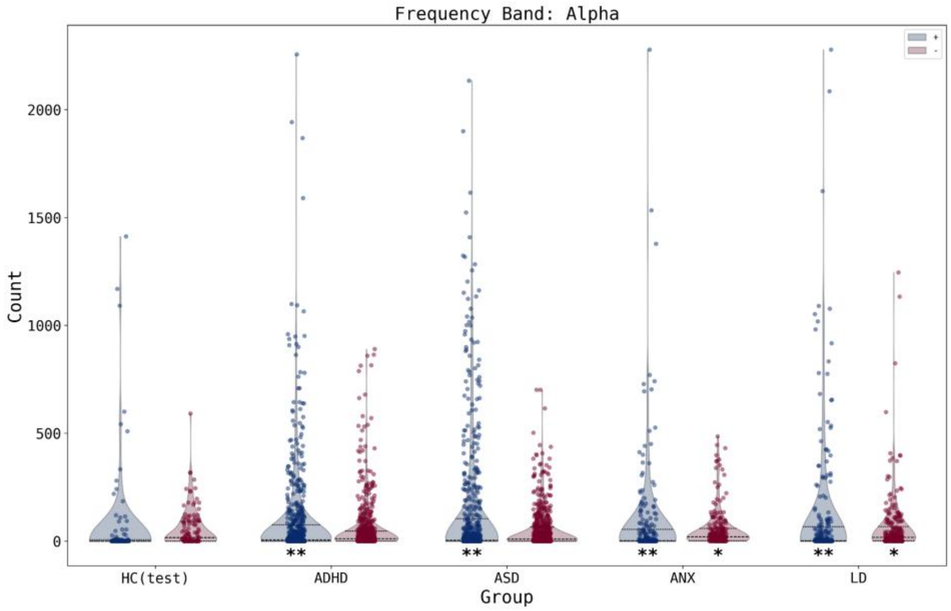


**Figure. S26 | Distribution of the number of extremely deviated connections per subject across groups in alpha band.** Blue (Red) violins represent positive (negative) deviations. (*) denotes significant difference between HC and cases. (*) denotes *p<0.05*, (**) denotes *p<0.01*.

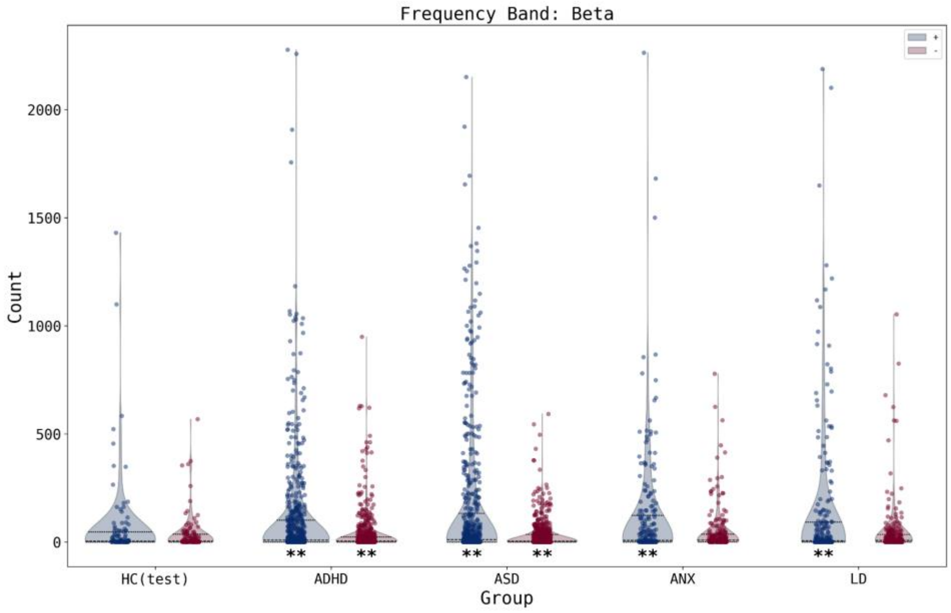


**Figure. S27 | Distribution of the number of extremely deviated connections per subject across groups in beta band.** Blue (Red) violins represent positive (negative) deviations. (*) denotes significant difference between HC and cases. (*) denotes *p<0.05*, (**) denotes *p<0.01*.

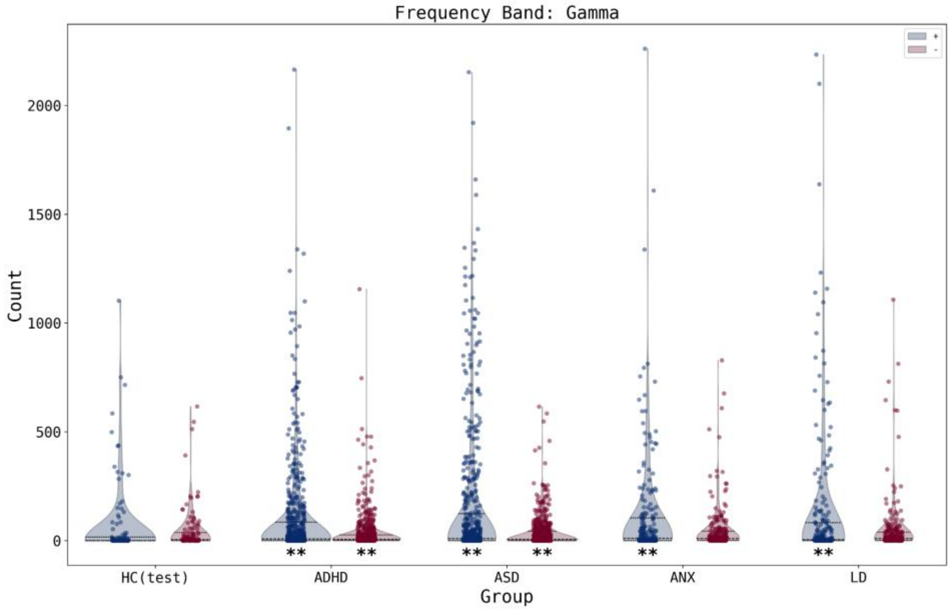


**Figure. S28 | Distribution of the number of extremely deviated connections per subject across groups in gamma band.** Blue (Red) violins represent positive (negative) deviations. (*) denotes significant difference between HC and cases. (*) denotes *p<0.05*, (**) denotes *p<0.01*.

**Table. S6 | Percentage of subjects exhibiting at least one extremely deviant channel and the median number of extreme deviations across groups in the delta frequency band.**

| Group (Delta) | % at least one positive deviation | Median [range] positive deviation | % at least one negative deviation | Median [range] negative deviation |
|---|---|---|---|---|
| HC(test) | 27.68 | 0.0 [0-104] | 23.21 | 0.0 [0-123] |
| ADHD | 32.31 | 0.0 [0-99] | 15.08 | 0.0 [0-124] |
| ASD | 41.15 | 0.0 [0-112] | 21.53 | 0.0 [0-121] |
| ANX | 28.24 | 0.0 [0-124] | 10.19 | 0.0 [0-59] |
| LD | 28.88 | 0.0 [0-124] | 19.4 | 0.0 [0-124] |

**Table. S7 | Percentage of subjects exhibiting at least one extremely deviant channel and the median number of extreme deviations across groups in the theta frequency band.**

| Group (Theta) | % at least one positive deviation | Median [range] positive deviation | % at least one negative deviation | Median [range] negative deviation |
|---|---|---|---|---|
| HC(test) | 31.25 | 0.0 [0-56] | 28.57 | 0.0 [0-124] |
| ADHD | 31.85 | 0.0 [0-123] | 27.54 | 0.0 [0-124] |
| ASD | 19.62 | 0.0 [0-124] | 43.75 | 0.0 [0-102] |
| ANX | 24.07 | 0.0 [0-114] | 18.52 | 0.0 [0-124] |
| LD | 25.86 | 0.0 [0-123] | 24.14 | 0.0 [0-124] |

**Table. S8 | Percentage of subjects exhibiting at least one extremely deviant channel and the median number of extreme deviations across groups in the alpha frequency band.**

| Group (Alpha) | % at least one positive deviation | Median [range] positive deviation | % at least one negative deviation | Median [range] negative deviation |
|---|---|---|---|---|
| HC(test) | 17.86 | 0.0 [0-115] | 27.68 | 0.0 [0-122] |
| ADHD | 17.08 | 0.0 [0-120] | 28.62 | 0.0 [0-124] |
| ASD | 11.46 | 0.0 [0-122] | 39.24 | 0.0 [0-111] |
| ANX | 13.89 | 0.0 [0-104] | 23.15 | 0.0 [0-124] |
| LD | 10.78 | 0.0 [0-62] | 24.57 | 0.0 [0-124] |

**Table. S9 | Percentage of subjects exhibiting at least one extremely deviant channel and the median number of extreme deviations across groups in the beta frequency band.**

| Group (Beta) | % at least one positive deviation | Median [range] positive deviation | % at least one negative deviation | Median [range] negative deviation |
|---|---|---|---|---|
| HC(test) | 22.32 | 0.0 [0-122] | 34.82 | 0.0 [0-110] |
| ADHD | 13.54 | 0.0 [0-124] | 25.23 | 0.0 [0-73] |
| ASD | 36.28 | 0.0 [0-124] | 42.01 | 0.0 [0-117] |
| ANX | 6.94 | 0.0 [0-61] | 19.44 | 0.0 [0-123] |
| LD | 15.52 | 0.0 [0-117] | 21.55 | 0.0 [0-123] |

**Table. S10 | Percentage of subjects exhibiting at least one extremely deviant channel and the median number of extreme deviations across groups in the gamma frequency band.**

| Group (Gamma) | % at least one positive deviation | Median [range] positive deviation | % at least one negative deviation | Median [range] negative deviation |
|---|---|---|---|---|
| HC(test) | 29.46 | 0.0 [0-66] | 26.79 | 0.0 [0-110] |
| ADHD | 20.62 | 0.0 [0-124] | 21.54 | 0.0 [0-123] |
| ASD | 23.61 | 0.0 [0-102] | 35.42 | 0.0 [0-115] |
| ANX | 25.93 | 0.0 [0-99] | 19.91 | 0.0 [0-119] |
| LD | 29.74 | 0.0 [0-124] | 20.26 | 0.0 [0-124] |

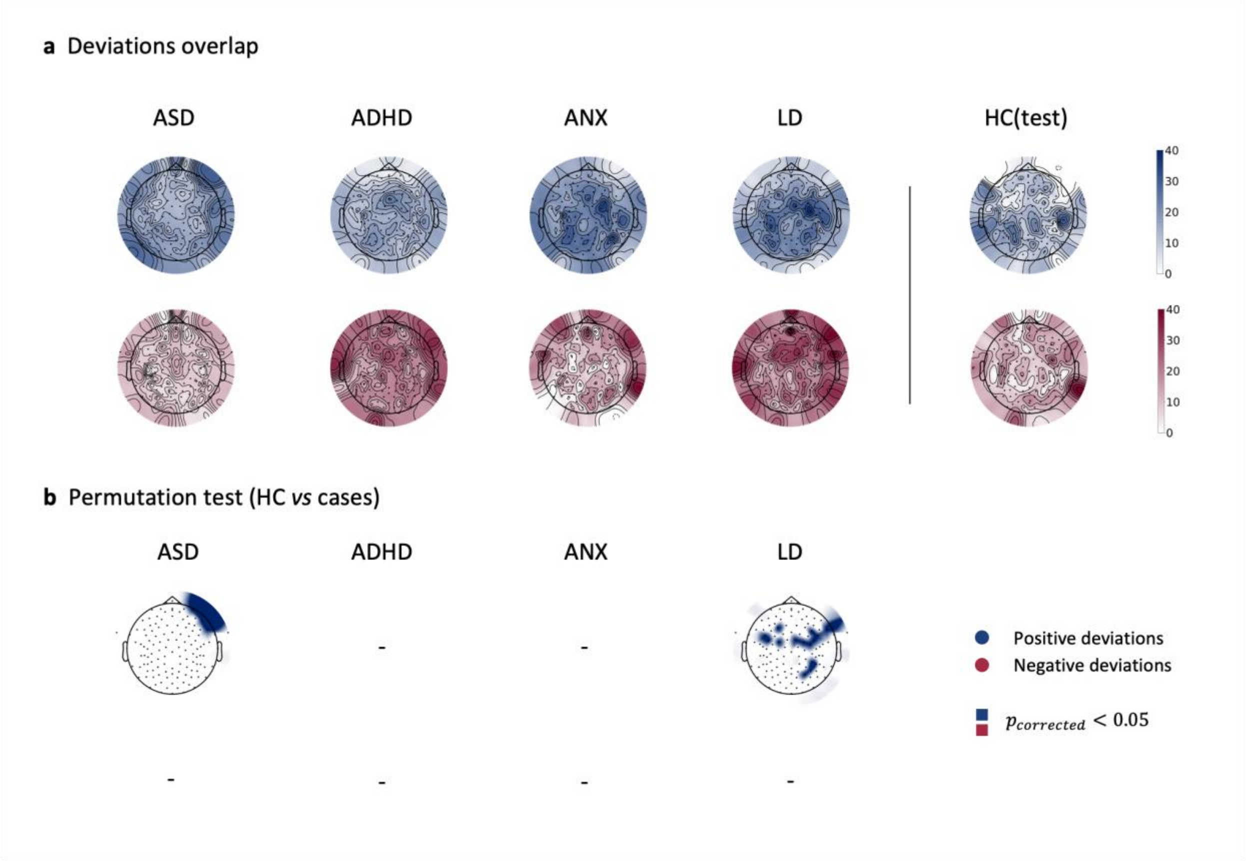


**Figure. S29 | Spectral features heterogeneity in delta band.** (a) Overlap maps of deviation scores for clinical groups and the held-out healthy control group (HC(test)), illustrating areas of common deviation. (b) channels showing significant differences between HC(test) and clinical groups, determined through group-based permutation tests (p<0.05, FDR corrected).

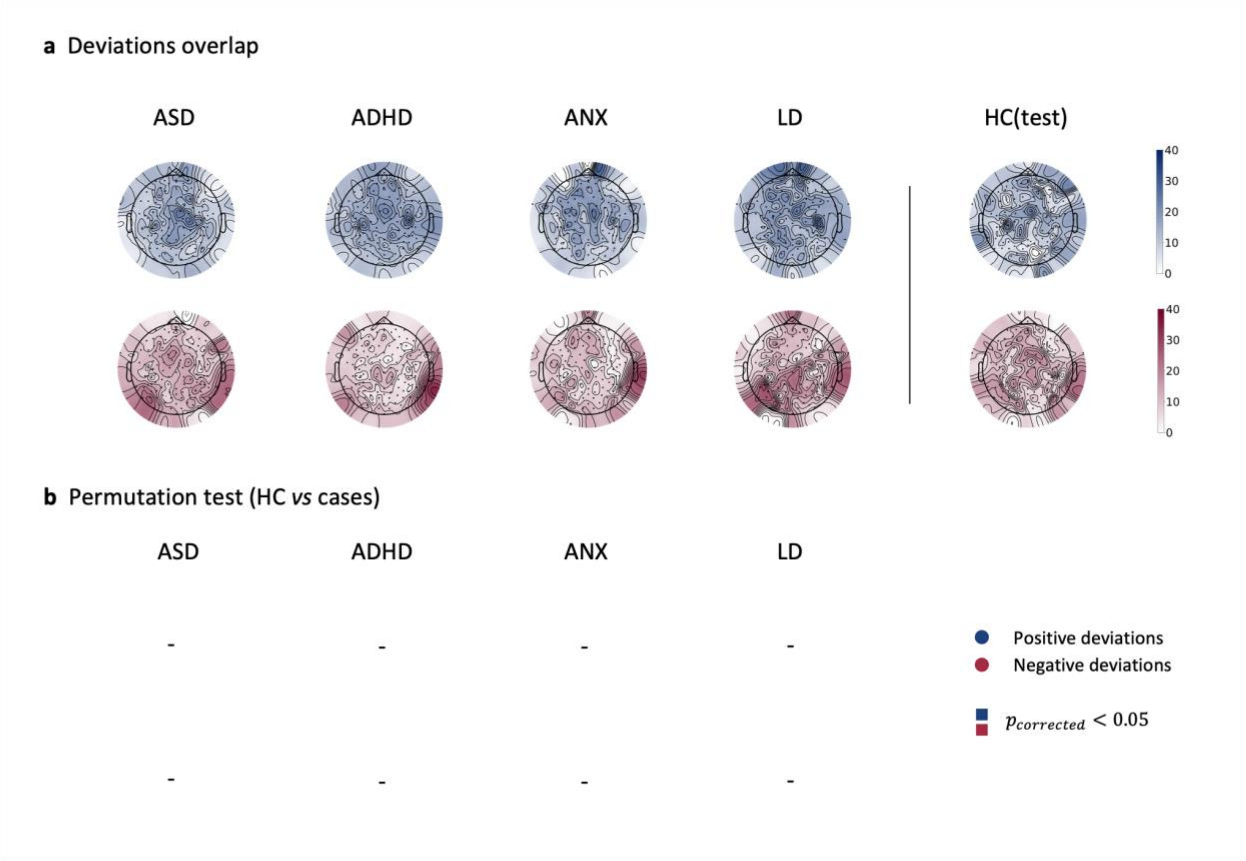


**Figure. S30 | Spectral features heterogeneity in theta band.** (a) Overlap maps of deviation scores for clinical groups and the held-out healthy control group (HC(test)), illustrating areas of common deviation. (b) channels showing significant differences between HC(test) and clinical groups, determined through group-based permutation tests (p<0.05, FDR corrected).

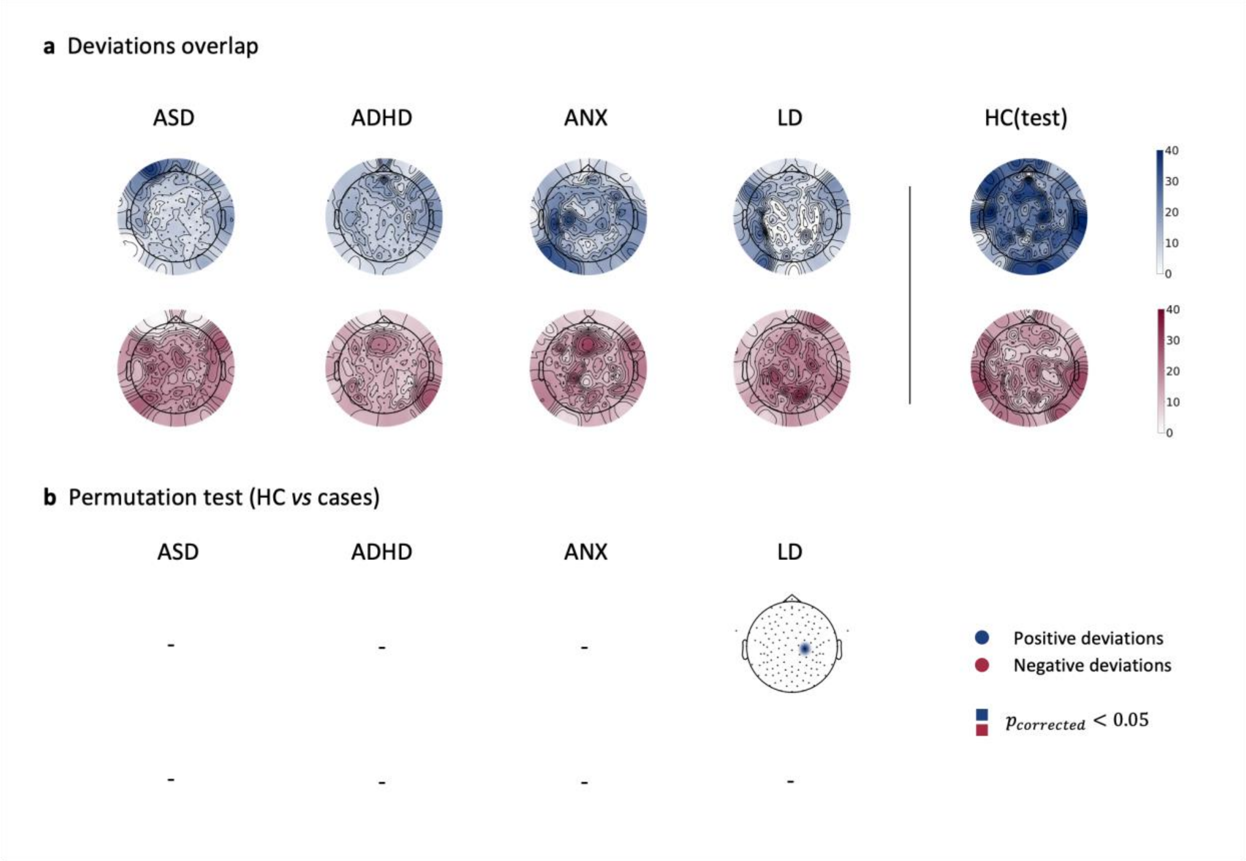


**Figure. S31 | Spectral features heterogeneity in alpha band.** (a) Overlap maps of deviation scores for clinical groups and the held-out healthy control group (HC(test)), illustrating areas of common deviation. (b) channels showing significant differences between HC(test) and clinical groups, determined through group-based permutation tests (p<0.05, FDR corrected).

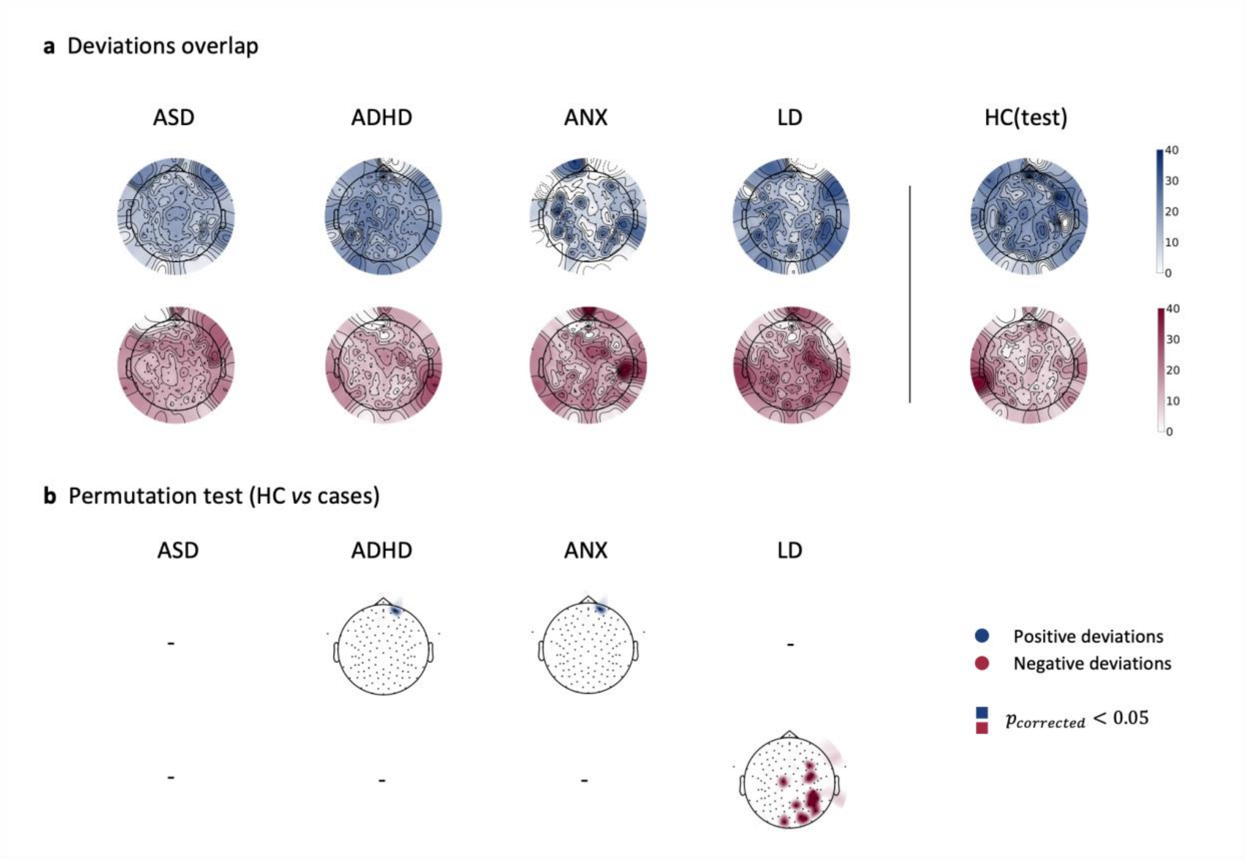


**Figure. S32 | Spectral features heterogeneity in beta band.** (a) Overlap maps of deviation scores for clinical groups and the held-out healthy control group (HC(test)), illustrating areas of common deviation. (b) channels showing significant differences between HC(test) and clinical groups, determined through group-based permutation tests (p<0.05, FDR corrected).

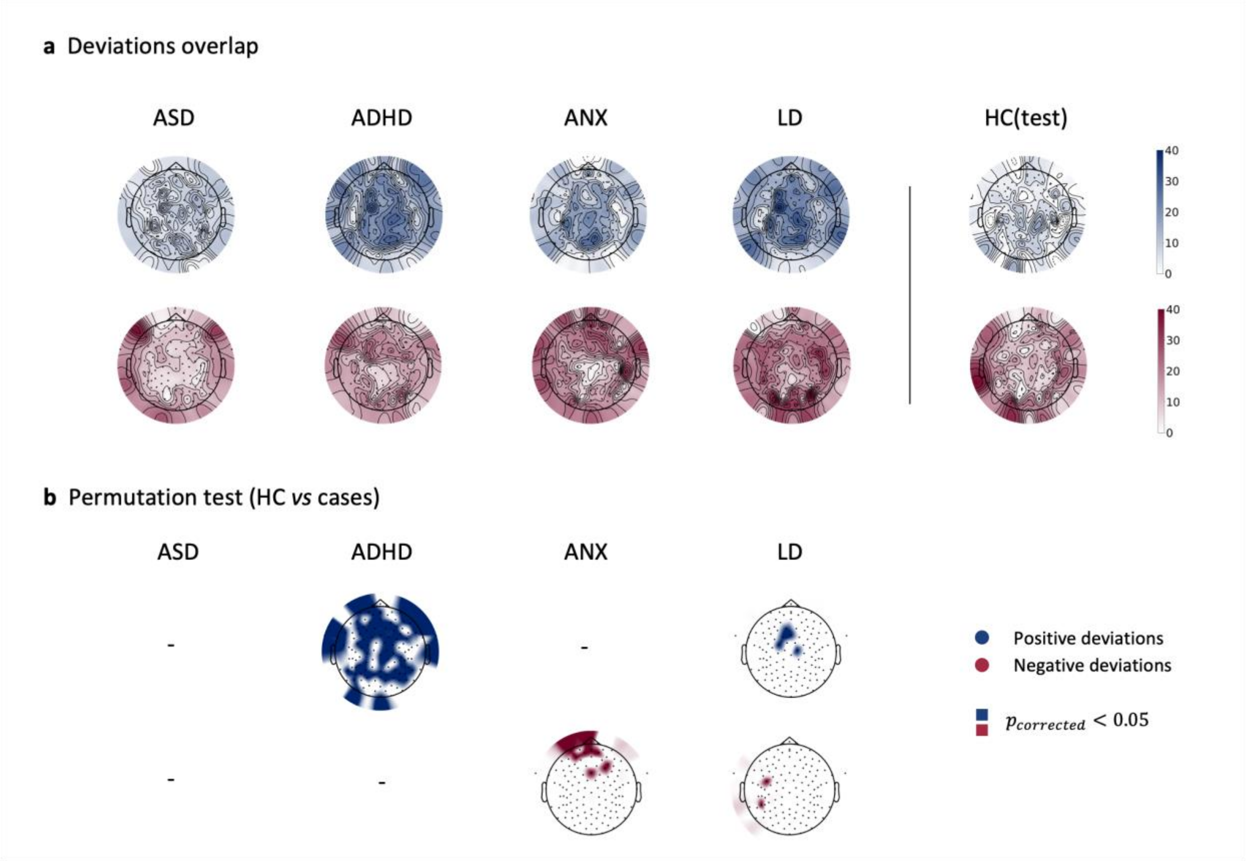


**Figure. S33 | Spectral features heterogeneity in gamma band.** (a) Overlap maps of deviation scores for clinical groups and the held-out healthy control group (HC(test)), illustrating areas of common deviation. (b) channels showing significant differences between HC(test) and clinical groups, determined through group-based permutation tests (p<0.05, FDR corrected).

**Table. S11 | Percentage of subjects exhibiting at least one extremely deviant connection and the median number of extreme deviations across groups in the delta frequency band.**

| Group (Delta) | % at least one positive deviation | Median [range] positive deviation | % at least one negative deviation | Median [range] negative deviation |
|---|---|---|---|---|
| HC(test) | 43.75 | 0.0 [0-1235] | 86.61 | 27.0 [0-604] |
| ADHD | 63.54 | 6.0 [0-2278] | 80.31 | 12.0 [0-877] |
| ASD | 70.66 | 7.5 [0-2144] | 70.66 | 11.0 [0-1050] |
| ANX | 63.26 | 5.0 [0-1679] | 81.4 | 18.0 [0-506] |
| LD | 57.33 | 4.0 [0-2278] | 81.47 | 15.5 [0-1321] |

**Table. S12 | Percentage of subjects exhibiting at least one extremely deviant connection and the median number of extreme deviations across groups in the theta frequency band.**

| Group (Theta) | % at least one positive deviation | Median [range] positive deviation | % at least one negative deviation | Median [range] negative deviation |
|---|---|---|---|---|
| HC(test) | 46.02 | 0.0 [0-449] | 74.34 | 9.0 [0-378] |
| ADHD | 62.77 | 4.0 [0-2263] | 81.85 | 10.0 [0-870] |
| ASD | 69.44 | 6.0 [0-2136] | 70.31 | 11.0 [0-1070] |
| ANX | 60.47 | 3.0 [0-2278] | 82.79 | 16.0 [0-412] |
| LD | 56.47 | 2.0 [0-2278] | 81.9 | 16.0 [0-1325] |

**Table. S13 | Percentage of subjects exhibiting at least one extremely deviant connection and the median number of extreme deviations across groups in the alpha frequency band.**

| Group (Alpha) | % at least one positive deviation | Median [range] positive deviation | % at least one negative deviation | Median [range] negative deviation |
|---|---|---|---|---|
| HC(test) | 37.72 | 0.0 [0-1412] | 68.42 | 16.5 [0-591] |
| ADHD | 63.23 | 6.0 [0-2256] | 80 | 11.5 [0-890] |
| ASD | 61.63 | 4.0 [0-2134] | 70.49 | 10.0 [0-701] |
| ANX | 56.54 | 2.5 [0-2278] | 81.78 | 20.5 [0-485] |
| LD | 55.6 | 2.0 [0-2278] | 81.47 | 18.0 [0-1245] |

**Table. S14 | Percentage of subjects exhibiting at least one extremely deviant connection and the median number of extreme deviations across groups in the beta frequency band.**

| Group (Beta) | % at least one positive deviation | Median [range] positive deviation | % at least one negative deviation | Median [range] negative deviation |
|---|---|---|---|---|
| HC(test) | 64.6 | 4.0 [0-1430] | 64.6 | 5.0 [0-568] |
| ADHD | 67.69 | 9.0 [0-2277] | 72.92 | 5.0 [0-949] |
| ASD | 72.74 | 12.0 [0-2151] | 66.32 | 4.0 [0-592] |
| ANX | 64.49 | 8.0 [0-2263] | 76.17 | 9.0 [0-778] |
| LD | 61.21 | 5.5 [0-2188] | 74.57 | 7.5 [0-1053] |

**Table. S15 | Percentage of subjects exhibiting at least one extremely deviant connection and the median number of extreme deviations across groups in the gamma frequency band.**

| Group (Gamma) | % at least one positive deviation | Median [range] positive deviation | % at least one negative deviation | Median [range] negative deviation |
|---|---|---|---|---|
| HC(test) | 46.02 | 0.0 [0-1102] | 65.49 | 5.0 [0-616] |
| ADHD | 67.23 | 8.0 [0-2164] | 74.92 | 6.0 [0-1155] |
| ASD | 71.35 | 10.0 [0-2152] | 65.97 | 6.0 [0-615] |
| ANX | 62.04 | 10.5 [0-2260] | 76.85 | 10.0 [0-828] |
| LD | 58.62 | 4.5 [0-2233] | 75 | 10.0 [0-1107] |
| | 43.75 | 0.0 [0-1235] | 86.61 | 27.0 [0-604] |

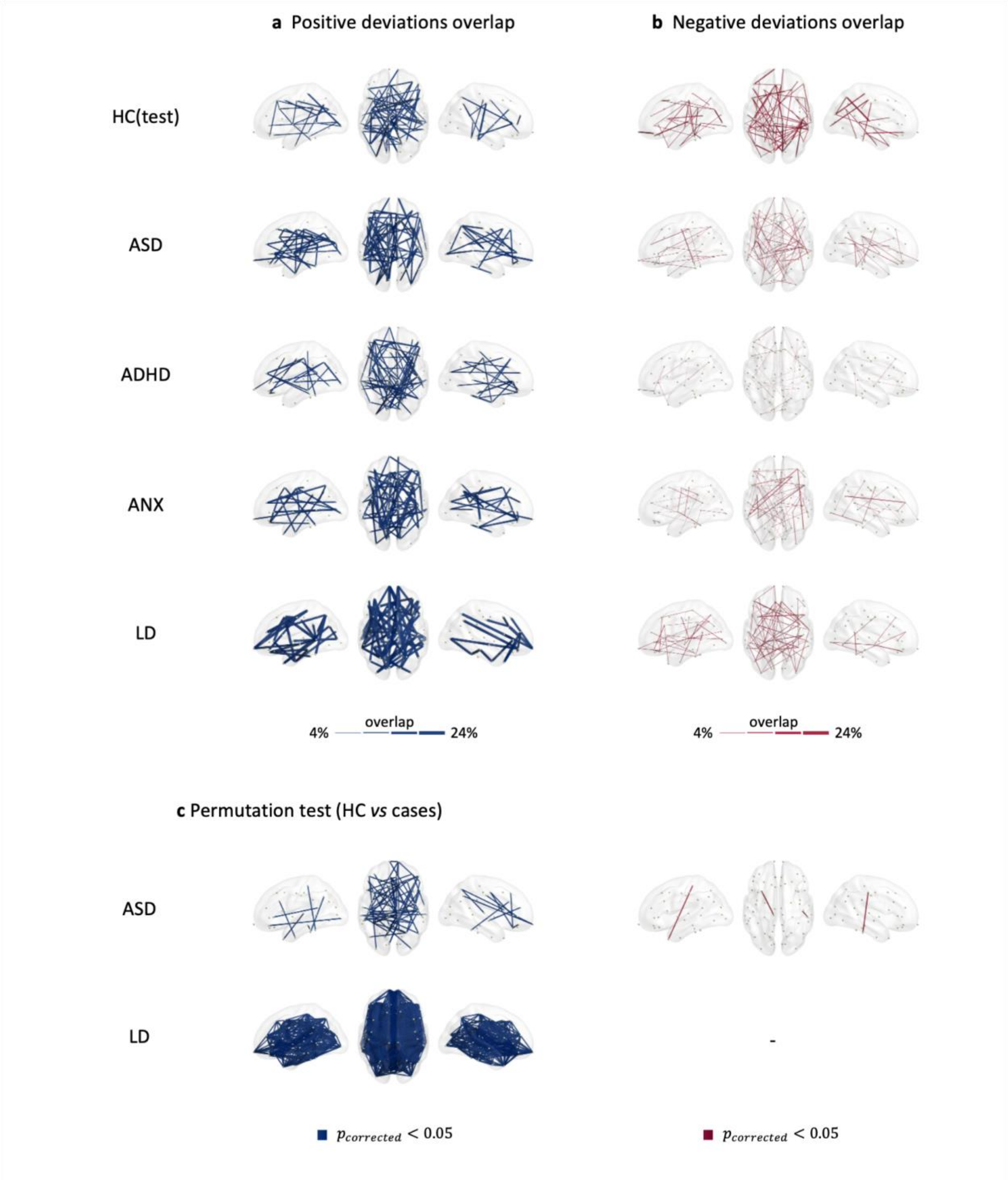


**Figure. S34 | Functional connectivity heterogeneity in delta band.** Overlap maps of (a) positive and (b) negative deviation scores for clinical groups and the held-out healthy control group (HC(test)) within the different frequency bands, illustrating areas of common deviation among patients (with only the highest 3% overlap values being plotted for visualization purposes). (c-d) functional connections showing significant differences between HC(test) and clinical groups at the different frequency bands, determined through group-based permutation tests (p<0.05, FDR corrected).

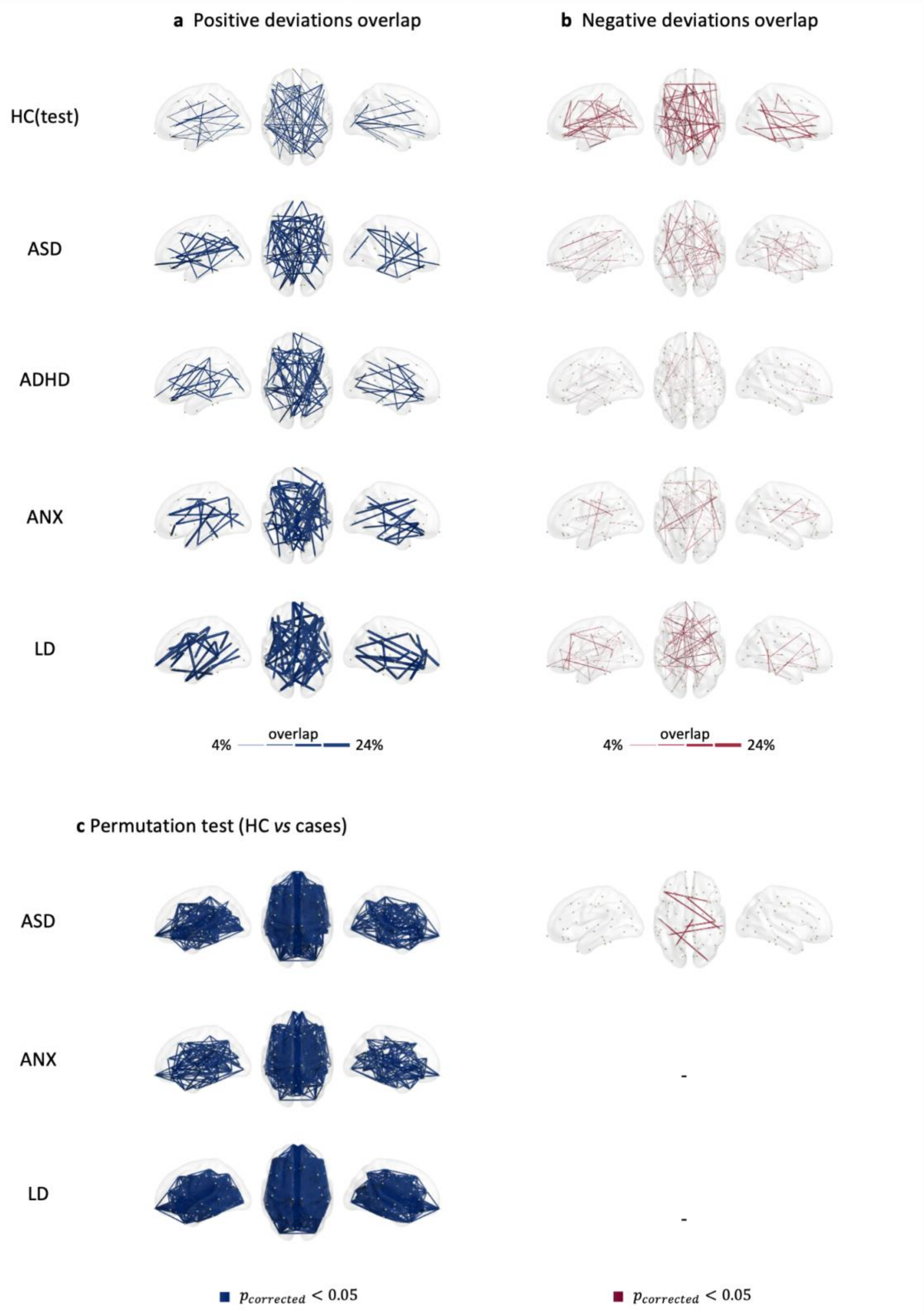

a Positive deviations overlap
b Negative deviations overlap
HC(test)
ASD
ADHD
ANX
LD
4%
overlap
24%
4%
overlap
24%
c Permutation test (HC vs cases)
ASD
ANX
LD
-
-
$p_{corrected} < 0.05$
$p_{corrected} < 0.05$

**Figure. S35 | Functional connectivity heterogeneity in theta band.** Overlap maps of (a) positive and (b) negative deviation scores for clinical groups and the held-out healthy control group (HC(test)) within the different frequency bands, illustrating areas of common deviation among patients (with only the highest 3% overlap values being plotted for visualization purposes). (c-d) functional connections showing significant differences between HC(test) and clinical groups at the different frequency bands, determined through group-based permutation tests ($p<0.05$, FDR corrected).

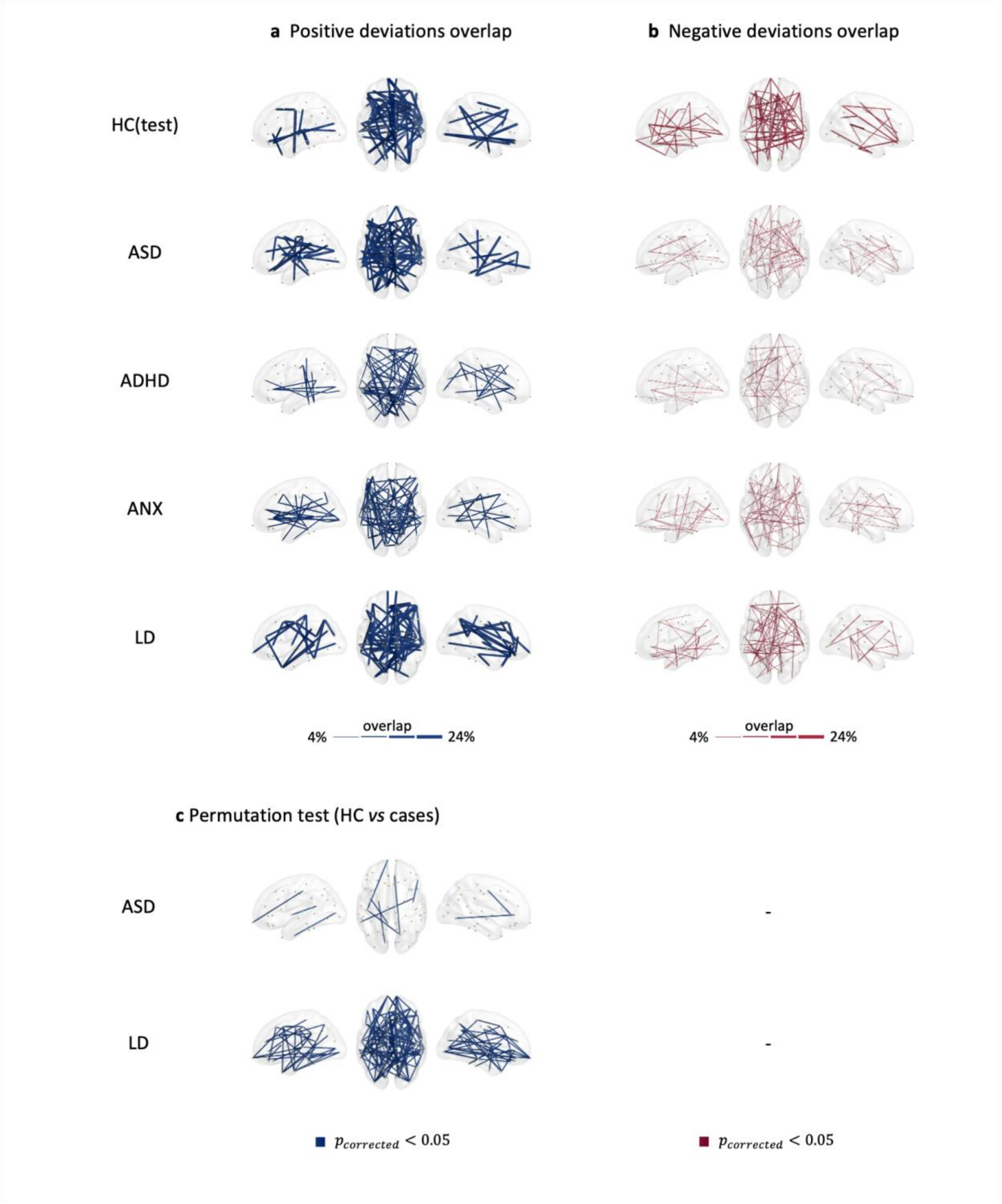


**Figure. S36 | Functional connectivity heterogeneity in alpha band.** Overlap maps of (a) positive and (b) negative deviation scores for clinical groups and the held-out healthy control group (HC(test)) within the different frequency bands, illustrating areas of common deviation among patients (with only the highest 3% overlap values being plotted for visualization purposes). (c-d) functional connections showing significant differences between HC(test) and clinical groups at the different frequency bands, determined through group-based permutation tests (p<0.05, FDR corrected).

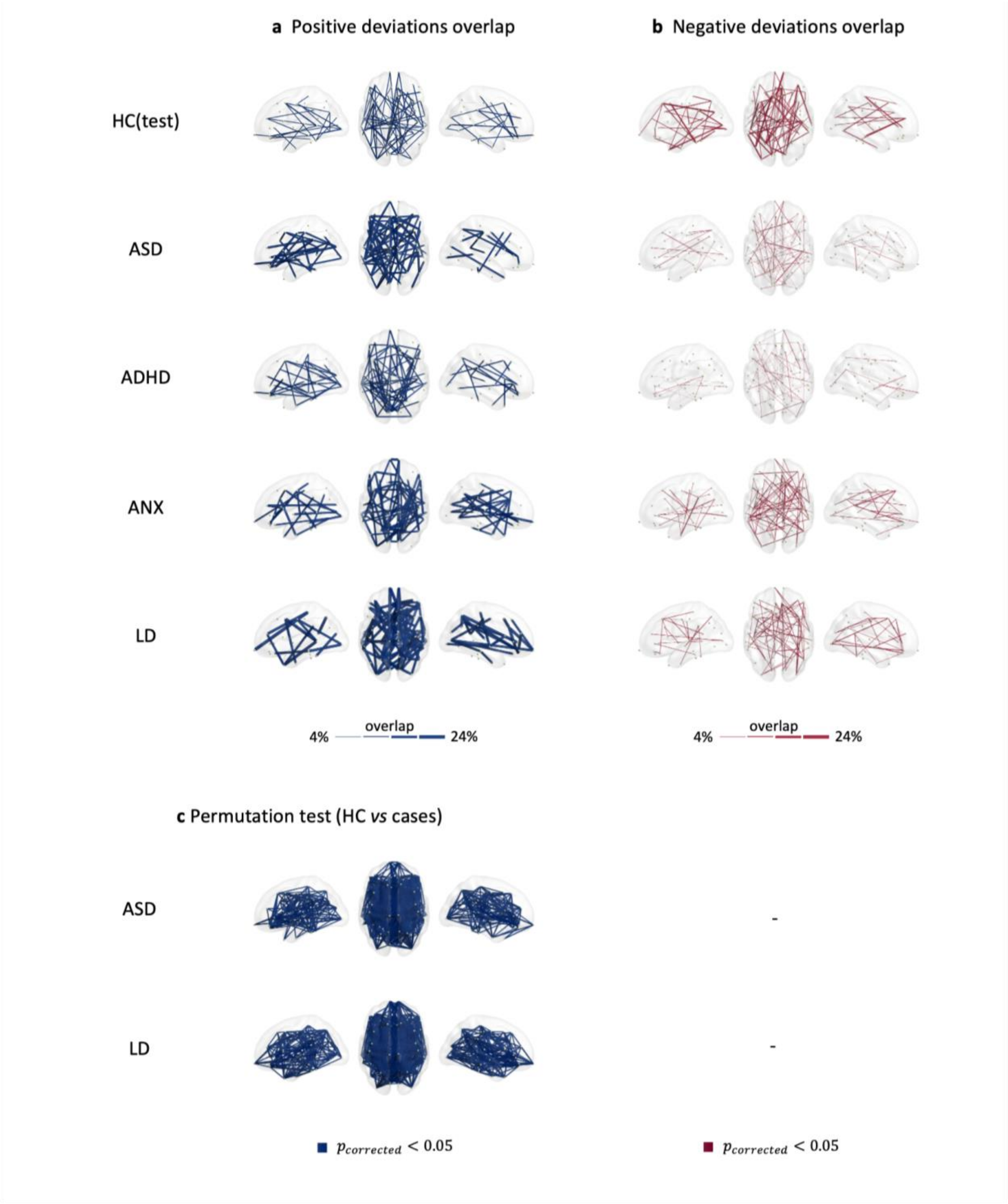

**Figure. S37 | Functional connectivity heterogeneity in beta band.** Overlap maps of (a) positive and (b) negative deviation scores for clinical groups and the held-out healthy control group (HC(test)) within the different frequency bands, illustrating areas of common deviation among patients (with only the highest 3% overlap values being plotted for visualization purposes). (c-d) functional connections showing significant differences between HC(test) and clinical groups at the different frequency bands, determined through group-based permutation tests ($p<0.05$, FDR corrected).

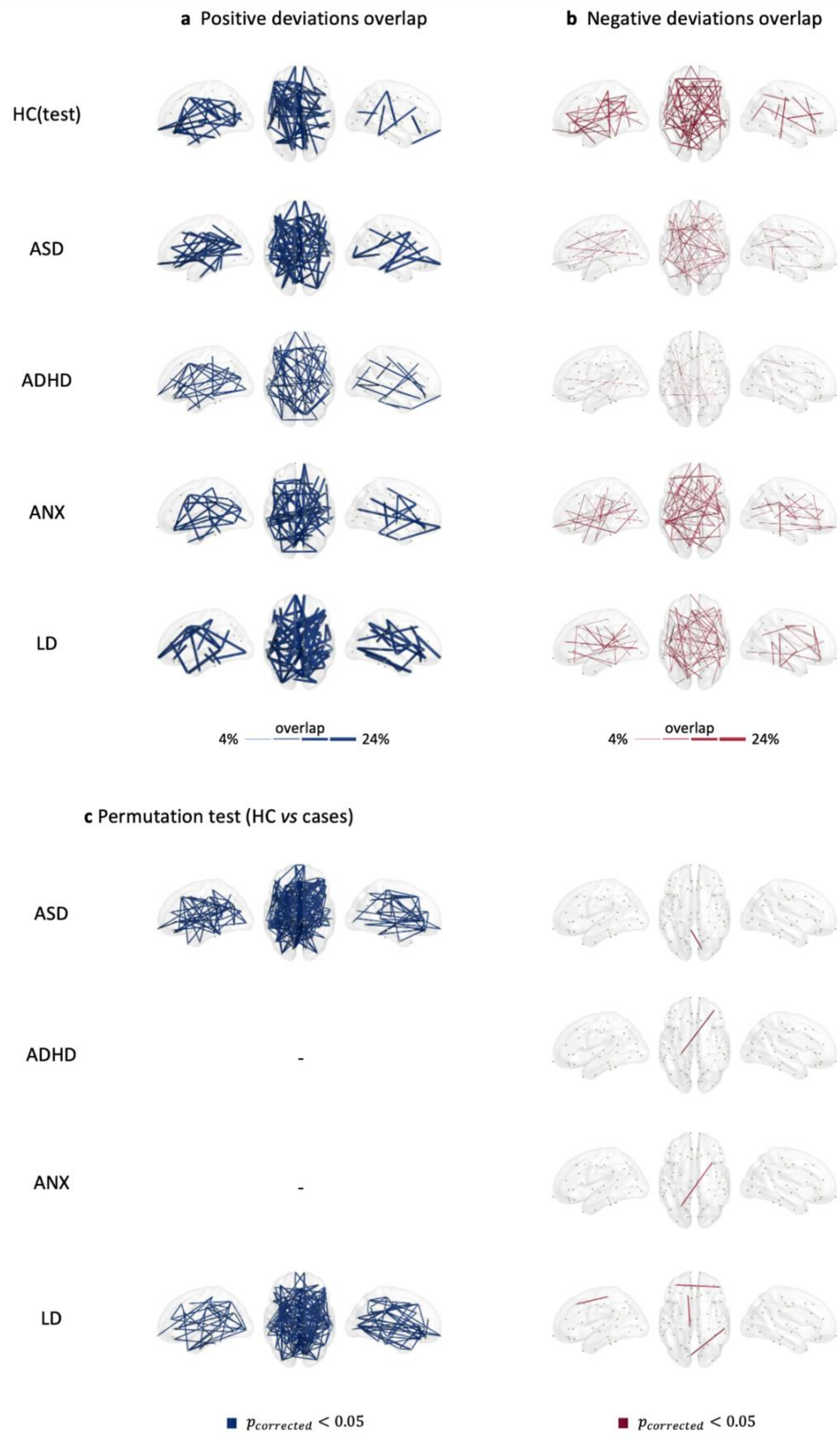
a Positive deviations overlap
b Negative deviations overlap
HC(test)
ASD
ADHD
ANX
LD
4%
overlap
24%
4%
overlap
24%
c Permutation test (HC vs cases)
ASD
ADHD
-
ANX
-
LD
$p_{corrected} < 0.05$
$p_{corrected} < 0.05$

**Figure. S38 | Functional connectivity heterogeneity in gamma band.** Overlap maps of (a) positive and (b) negative deviation scores for clinical groups and the held-out healthy control group (HC(test)) within the different frequency bands, illustrating areas of common deviation among patients (with only the highest 3% overlap values being plotted for visualization purposes). (c-d) functional connections showing significant differences between HC(test) and clinical groups at the different frequency bands, determined through group-based permutation tests ($p<0.05$, FDR corrected).

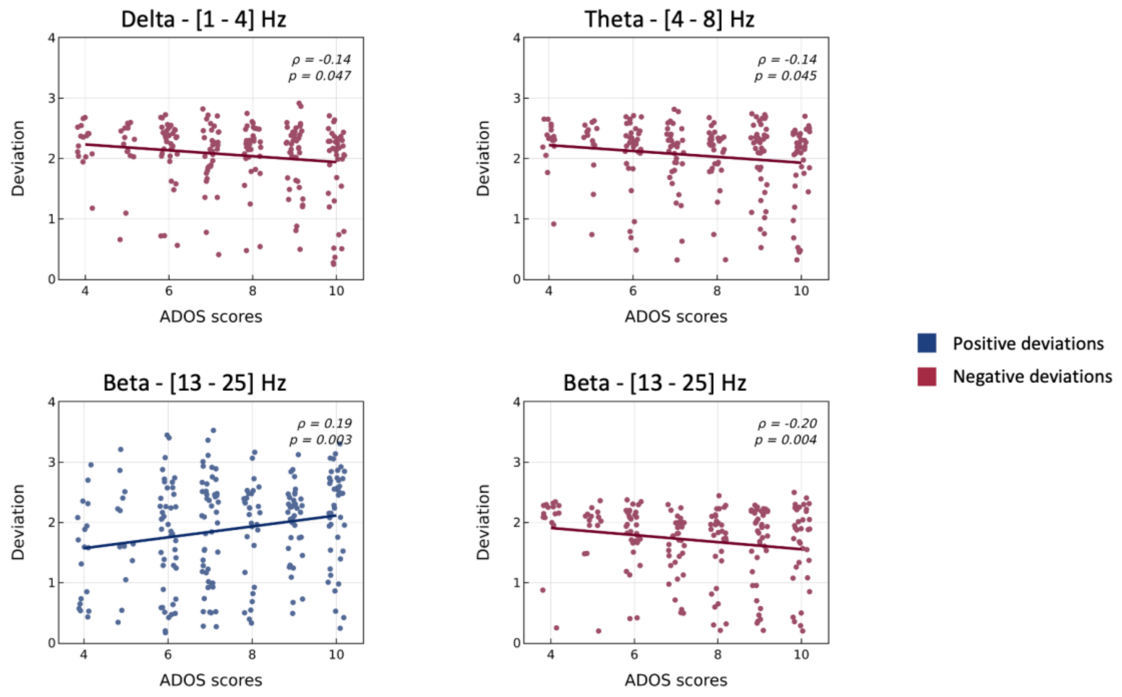


**Figure. S39 | Correlation between ASD subjects' global deviation scores and clinical assessment scores (ADOS) in delta, theta, and beta bands.**

Lord, Catherine, M. Rutter, P. DiLavore, S. Risi, K. Gotham, S. Bishop, and Others. 2012. “Autism Diagnostic Observation Schedule--2nd Edition (ADOS-2).” *Los Angeles, CA: Western Psychological Corporation* 284.

McPartland, James C., Raphael A. Bernier, Shafali S. Jeste, Geraldine Dawson, Charles A. Nelson, Katarzyna Chawarska, Rachel Earl, et al. 2020. “The Autism Biomarkers Consortium for Clinical Trials (ABC-CT): Scientific Context, Study Design, and Progress Toward Biomarker Qualification.” *Frontiers in Integrative Neuroscience* 14 (April): 16.

Pelphrey, Kevin. 2014. “Multimodal Developmental Neurogenetics of Females with ASD.”
